# A CPC-shelterin-BTR axis regulates mitotic telomere deprotection

**DOI:** 10.1101/2024.01.09.574754

**Authors:** Diana Romero-Zamora, Samuel Rogers, Ronnie Ren Jie Low, Andrew B. Robinson, Scott G. Page, Blake J. E. Lane, Noa Lamm, Fuyuki Ishikawa, Makoto T. Hayashi, Anthony J. Cesare

**Affiliations:** Graduate School of Biostudies, Kyoto University, Yoshida-Konoe, Sakyo, Kyoto, 606-8501 Japan; IFOM-KU Joint Research Laboratory, Graduate School of Medicine, Kyoto University, Yoshida-Konoe, Sakyo, Kyoto, 606-8501 Japan; Children’s Medical Research Institute, University of Sydney, Sydney, New South Wales 2145, Australia; IFOM ETS, The AIRC Institute of Molecular Oncology, Via Adamello 16, 20139 Milan, Italy

**Keywords:** Telomeres, mitosis, shelterin, TRF1, TRF2, chromosome passenger complex, AURKB, BLM-TOP3A-RMI1/2, mitotic telomere deprotection, t-loop

## Abstract

Telomeres prevent ATM activation by sequestering chromosome termini within telomere loops (t-loops). Mitotic arrest promotes telomere linearity and a localized ATM-dependent telomere DNA damage response (DDR) through an unknown mechanism. Using unbiased interactomics, biochemical screening, molecular biology, and super-resolution imaging, we found that mitotic arrest-dependent (MAD) telomere deprotection requires the combined activities of the Chromosome passenger complex (CPC) on shelterin, and the BLM-TOP3A-RMI1/2 (BTR) complex on t-loops. During mitotic arrest, the CPC component Aurora Kinase B (AURKB) phosphorylated both the TRF1 hinge and TRF2 basic domains. The former enhanced CPC and TRF1 interaction through the CPC Survivin subunit, while the latter promoted telomere linearity, telomere DDR activation dependent upon BTR double Holliday junction dissolution activity, and mitotic death. We identify that the TRF2 basic domain functions in mitosis-specific telomere protection and reveal TRF1 regulation over a physiological ATM-dependent telomere DDR. The data demonstrate that MAD telomere deprotection is a sophisticated active mechanism that exposes telomere ends to signal mitotic stress.

## INTRODUCTION

Telomeres are the specialized nucleoprotein structures at eukaryotic chromosome termini. The paramount telomere activity is protection of chromosome ends through localized regulation of DDR and DNA repair activity by the telomere-specific shelterin protein complex^1^. Telomere deprotection refers to physiological outcomes where chromosome end protection is compromised through biological processes, typically in the presence of wild type shelterin^2^.

Evidence supports the telomere-specific DDR that defines mammalian telomere deprotection resulting from macromolecular changes in telomere structure^3–5^. Diverse eukaryotic species arrange their telomeres into the lariat t-loop configuration, where the terminal 3’-overhang of the G-rich telomere sequence strand invades the duplex telomere DNA in cis^6–10^. T-loops in somatic mammalian tissues require the TRFH domain of the TRF2 shelterin subunit^11,12^. Partial TRF2 depletion, or expression of TRF2 TRFH domain mutants in a *Trf2^−/−^* background, leads to a telomere-specific and ATM-dependent DDR^13,14^, corresponding with a transition from looped to linear telomeres^11^. Pluripotent murine embryonic stem cells (mESCs) devoid of TRF2 retain t-loops and suppress ATM activity at chromosome ends^15,16^. However, deletion of two shelterin genes in mESCs, *Trf1* and *Trf2*, confer linear telomeres and a corresponding ATM-dependent telomere DDR^15^. The collective data are consistent with a model where chromosome termini are sequestered within t-loops to prevent ATM activation by the naturally occurring DNA ends.

Canonical telomere deprotection, elicited through aging-dependent telomere erosion, progressively activates two independent tumour suppressive programs^2^. Replicative senescence occurs first and is mediated by a telomere-specific DDR that promotes p53-dependent proliferative arrest^4,17^. p53 compromised cells are refractory to this telomere DDR and continue cell division^17,18^ until excessive telomere erosion confers end-to-end chromosome fusions and cell death at crisis^5,19^. Cell lethality during crisis is mediated primarily by autophagic death signalled through the innate immune response^20,21^. A minor proportion of crisis cells, however, perish during mitosis in a MAD manner^22^. Similar mitotic death dependent upon mitotic arrest occurs in response to chemotherapeutic mitotic poisons or lethal replication stress^23,24^. These physiological mitotic death events are promoted in part through a poorly understood and non-canonical mechanism we term MAD-telomere deprotection^22,24–26^.

MAD-telomere deprotection is a unique phenomenon where telomere-independent stress promotes active telomere deprotection. In addition to potentiating mitotic death^22^, MAD-telomere deprotection also promotes p53-dependent proliferative arrest in the following interphase should cells escape mitotic arrest^25,27^. Co-localization of telomeres and DDR markers are termed telomere deprotection induced foci (TIF)^28^. When these events occur due to mitotic arrest, we term them MAD-TIF. MAD-TIF arise in human cells following four or more hours of mitotic arrest and increase in number as mitotic delay persists^25^. Diverse genetic or pharmacological interventions that arrest mitosis all confer MAD-TIF^22,24,25^, indicating that MAD-telomere deprotection is a physiological response to general mitotic delay. Additionally, MAD-TIF occur irrespective of telomere length and arise independent of telomerase activity^25^. The DDR observed with MAD-TIF is ATM-dependent, suppressed by TRF2 over-expression, and correlates with a transition from looped to linear telomeres^11,25,29^. This suggests that MAD-TIF result from t-loop opening during mitotic arrest. Congruent with a regulated phenomenon, MAD-TIF require AURKB activity but not the mitotic kinases Aurora A or MPS1^25^.

MAD-telomere deprotection presents a unique opportunity to explore critical open questions in telomere protection. Foremost is, does shelterin participate in the active process of MAD-telomere deprotection? If so, this changes the conceptual shelterin paradigm from a solely protective complex to a dynamic entity capable of transitioning telomeres between protected and deprotected states. Furthermore, shelterin components in somatic cells are largely considered to regulate individual DDR activities, with TRF2 being the sole mediator of ATM suppression^1,30^. If additional shelterin components regulate ATM-dependent MAD-TIF, this would represent an unexpected expansion of shelterin protective activities. Additionally, while t-loop junctions are typically presented as a 3-way displacement loop (D-loop), the exact molecular structure at the t-loop insertion point remains conjecture. Alternative configurations include a 4-way Holliday junction (HJ), or a double Holliday junction (dHJ) observed during homologous recombination (HR). The N-terminal TRF2 basic domain binds three- and four-way DNA structures in a sequence independent manner and is proposed to protect t-loop junctions^31–34^. It remains unclear if the telomere protection afforded by the TRF2 basic domain is ubiquitous or cell cycle specific. Finally, how t-loop junctions are resolved during mitosis, and if this occurs spontaneously or through recruitment of an independent factor, also remains unknown.

Here we present the development of an unbiased telomere interactomics tool and its application to identify regulators of MAD-telomere deprotection. This revealed that the CPC and BTR complexes associate with TRF1 specifically during mitotic arrest. The CPC is a key mitotic regulator comprised of AURKB, INCENP, Survivin, and Borealin^35^, while BTR promotes dHJ dissolution^36^. We found that AURKB phosphorylated both the TRF1 hinge and the TRF2 basic domains during mitotic arrest, and that these modifications were requisite for BTR-dependent MAD-telomere deprotection. The data reveal unexpected regulation of ATM-dependent MAD-TIF by TRF1 and demonstrate that the TRF2 basic domain protects t-loops from BTR-dependent dHJ dissolution specifically during mitosis. Collectively, the data support a model where co-ordinated post-translational shelterin modifications promote t-loop unwinding during mitotic arrest to remove damaged cells from the cycling population.

## RESULTS

### TRF1 interacts with the CPC and BLM during mitotic arrest

To identify proteins that interact with human telomeres during mitotic arrest we utilized an unbiased and time-resolved proximity biotinylation strategy. When activated with biotin-phenol and 1 mM hydrogen peroxide, engineered ascorbic acid peroxidase (APEX2) promiscuously labels proteins within a 10 – 20 nm radius in 60 seconds^37^ (Fig S1A). We created tri-cistronic lentivectors harboring mCherry:P2A:PuroR:T2A:FLAG-APEX2 or FLAG-APEX2-TRF1 and transduced HeLa cultures. Cells were selected for Puromycin resistance and sorted on low mCherry expression to equalize FLAG-APEX2 and FLAG-APEX2-TRF1 expression between conditions (Fig 1A). We verified APEX2 activity after activation through streptavidin staining of whole cell extracts (Fig S1B). Cytological assessment of streptavidin staining in transduced cultures following one minute of APEX2 activation revealed diffuse localization in Flag-APEX2 cells and spatial enrichment at the telomeres in Flag-APEX2-TRF1 cultures (Fig. 1B). This was indicative of tight spatiotemporal Flag-APEX2-TRF1 labelling, allowing for unbiased time-resolved telomere interactomics.

**Figure 1.**
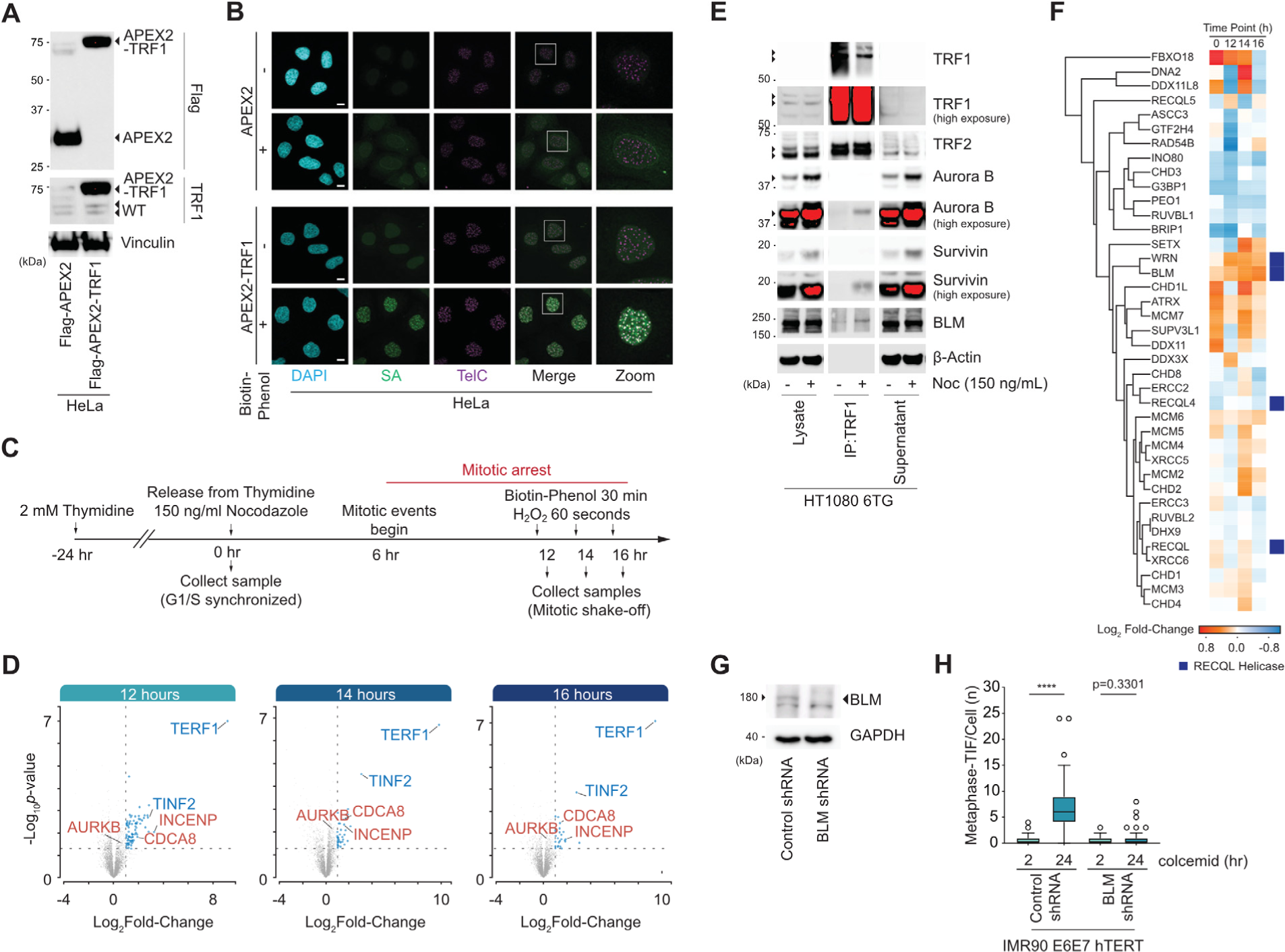
TRF1 interactomics reveal CPC and BLM function in MAD telomere deprotection. **A.** Immunoblots of whole cell extracts derived from Flag-APEX2 or Flag-APEX2-TRF1 expressing HeLa cells (representative example of n = 3 biological replicates). **B.** Micrographs of combined telomere fluorescence in situ hybridization (FISH, TelC) and streptavidin (SA) labelling as detected by immunofluorescence (IF) in Flag-APEX2 or Flag-APEX2-TRF1 expressing HeLa cells with or without APEX activation by biotin-phenol. (representative example of n = 3 biological replicates). **C.** Timeline of mitotic arrest enrichment and biotin labelling for the HeLa Flag-APEX2-TRF1 interactomics. **D.** Volcano plots of protein enrichment from Flag-APEX2-TRF1 interactomics described in (C) and Fig. S1C, D. Proteins enriched from Flag-APEX2-TRF1 samples were compared to respective Flag-APEX2 timepoints and plotted as a function of Log_2_ fold-change and –Log_10_ p-value, indicating enriched members of CPC (red) and Shelterin (blue) complexes (student’s T-test of n = 5 biological replicates). **E.** Immunoblots of endogenous TRF1 immuno-precipitates from HT1080 6TG cells. Where indicated, cells were treated for 16 hr with150 ng mL^−1^ Nocodazole (Noc) immediately prior to sample collection (representative example of n = 3 biological replicates). **F.** Hierarchical clustering of proteins classified by gene-ontology with DNA helicase activity (GO:0003678), that were detected by Flag-APEX2-TRF1 interactomics as described in (C). Colours indicate Log_2_-fold change comparing APEX2-TRF1 and APEX2 samples. RECQL helicases are identified by a blue box on the right. **G.** Immunoblots of whole cell extracts derived from IMR90 E6E7 hTERT fibroblasts expressing control or BLM shRNAs. Extracts were collected 5 days post shRNA transduction (representative example of n = 3 biological replicates). **H.** Metaphase-TIF in shRNA transduced IMR90 E6E7 hTERT fibroblasts treated with 2 or 24 hours of 100 ng mL^−1^ colcemid (n = 3 biological replicates of 15 or 30 metaphases per replicate for 2 hours and 24 hours colcemid, respectively, all data points are combined in a Tukey box plot, Mann-Whitney U test, ****p<0.0001).

To enrich for mitotically arrested cells, we synchronized Flag-APEX2 and Flag-APEX2-TRF1 expressing HeLa cells in G1/S using a single 2 mM thymidine block for 24 hours. HeLa cells enter mitosis 6 hours after thymidine washout^25^. We released G1/S synchronized cultures into media containing 150 ng mL^−1^ of the microtubule poison Nocodazole to arrest cells in the subsequent mitosis. APEX2 or APEX2-TRF1 were activated 12, 14, or 16 hours after release from G1/S, corresponding with 6, 8, and 10 hours of mitotic arrest (Fig 1C). Following APEX2 or APEX2-TRF1 activation for one minute, we collected samples and recovered biotinylated proteins from cell lysates via streptavidin pulldown (Fig. S1C). For these experiments, Flag-APEX2 samples provide reference for non-specific interacting factors, while G1/S synchronized (0-hour post-release) Flag-APEX2-TRF1 samples provide reference for proteins that ubiquitously associate with telomeres.

Streptavidin recovered material was analysed via LC-MS/MS. We refined 7,630 total identifications to 1,400 significant proteins, of which 103 were deemed potential Flag-APEX2-TRF1 specific interactors with a Log_2_Fold-Change > 1 at 12, 14, and, 16 hours post-release (Fig. S1D). The shelterin components TRF1 (TERF1) and TIN2 (TINF2) were identified as higher confidence interactors by more stringent statistical testing (Fig. 1D, and S1E). Recovery of TRF2 (TERF2) and other shelterin components by Flag-APEX2-TRF1 modestly decreased in abundance during mitotic arrest (Fig. S1E), consistent with prior observations of reduced TRF2 at mitotic telomeres during MAD-telomere deprotection^25,26^.

Pathway analysis of the 103 potential Flag-APEX2-TRF1 interactors identified an enrichment of mitotic regulators (Fig. S1F). This included the kinetochore localized proteins CASC5, SPDL1, CENPE, DSN1, and SGOL2^38^. Additionally, the CPC components AURKB, INCENP, and CDCA8 (Borealin), were strongly enriched within the Flag-APEX2-TRF1 samples (Fig. 1D and S1G). AURKB, INCENP and Borealin follow a similar trend in recovery relative to cell synchrony; all were largely absent at G1/S, 0 hours after release from the Thymidine block, and peaked at 12 hours post-release corresponding to early MAD-telomere deprotection (Fig. S1G). HT1080 6TG cells readily display MAD-telomere deprotection^24^. Co-immunoprecipitations (co-IP) of endogenous TRF1 from HT1080 6TG cells enriched for mitotic arrest through treatment with 150 ng mL^−1^ Nocodazole for 16 hours also recovered AURKB and the additional CPC component Survivin (Fig. 1E). Similar TRF1 and CPC interactions were not observed in asynchronous controls (Fig. 1E). TRF1 thus interacts with mitotic regulatory proteins critical for MAD-telomere deprotection^25^, specifically during mitotic arrest, in multiple human cell lines.

MAD-TIF correlate with a transition from looped to linear telomeres^11^, and t-loop unwinding is postulated to require helicase activity^39^. Analysis of our interactomics dataset revealed 39 proteins with DNA helicase activity (GO:0003678), 11 of which were significantly detected at the telomeres during mitotic arrest (Fig. 1F). Among these helicases, BLM is a known TRF1 interactor^40^ that was proposed to promote dissolution of telomere dHJs should they result from t-loop branch migration^32^. BLM was strongly co-immunoprecipitated with endogenous TRF1 in mitotically arrested but not asynchronous cultures (Fig. 1E). This piqued our interest in BLM as a potential regulator of MAD-telomere deprotection.

MAD-TIF are typically studied in human primary diploid IMR90 fibroblasts^25^. Here we used hTERT immortalized IMR90 that also express papilloma virus serotype 16 E6 and E7 (IMR90 E6E7 hTERT) to respectively inhibit p53 and RB. E6E7 expression facilitates cell division when shelterin proteins are targeted^29^. To measure metaphase-TIF, cells are cytocentrifuged onto class coverslips and stained with telomere fluorescent in situ hybridization (FISH) and immunoflourenece against γ-H2AX^41^. Assessment of MAD-TIF requires mitotic arrest. IMR90 E6E7 hTERT were treated for 2 or 24 hours with 100 ng mL^−1^ of the microtubule poison colcemid (Fig. 1G-H). With endogenous shelterin, colcemid treatment for two hours is insufficient to induce MAD-telomere deprotection^25^. Under these conditions, metaphase-TIF present after two hours of colcemid represent TIF inherited from the prior interphase^29^ (Fig. 1H). The difference in metaphase-TIF between 2 and 24 hours colcemid represent MAD-TIF induced through mitotic delay (Fig. 1H). We found that BLM shRNA depletion robustly suppressed MAD-TIF in the 24 hr colcemid treated cells. TRF1 therefore interacts with both AURKB and BLM during mitotic arrest, and BLM promotes MAD telomere deprotection.

### AURKB phosphorylates TRF1 to promote MAD telomere deprotection

The above results inferred a previously unidentified role for TRF1 in MAD-telomere deprotection. To explore further, we shRNA depleted endogenous TRF1 in IMR90 E6E7 hTERT and complemented with exogenous shRNA resistant TRF1 alleles (TRF1^shR^)(Fig 2A, B, and S2A-E). To our surprise, TRF1 depletion suppressed MAD-TIF in cultures treated with colcemid for 24 hours (Fig 2B and S2A). MAD-TIF were rescued with TRF1^shR^-wild type (WT) but not TRF1^shR^-ΔBLM which carries a deletion of the BLM-binding motif^40^ (Fig. 2B and S2B). In silico analysis of the TRF1 protein sequence identified three potential (R/K)_1-3_-X-S/T AURKB phosphorylation sites^42^, Ser296, Ser354, and Thr358, in the hinge domain near the BLM-binding motif. Ectopic expression of TRF1^shR^ alleles carrying an Ala mutation of all or each residue revealed that Ser354 and Thr358 were required for MAD-TIF formation (Fig. 2B, and S2C, D). Mutation of both S354 and T358 to phospho-mimetic Asp (TRF1^shR^-2D) restored MAD-TIF (Fig. 2C, and S2E), suggestive of TRF1 phospho-regulation being requisite for MAD-telomere deprotection. We note that S354 is absent from the TRF1^shR^-ΔBLM allele that fails to rescue MAD-TIF.

**Figure 2.**
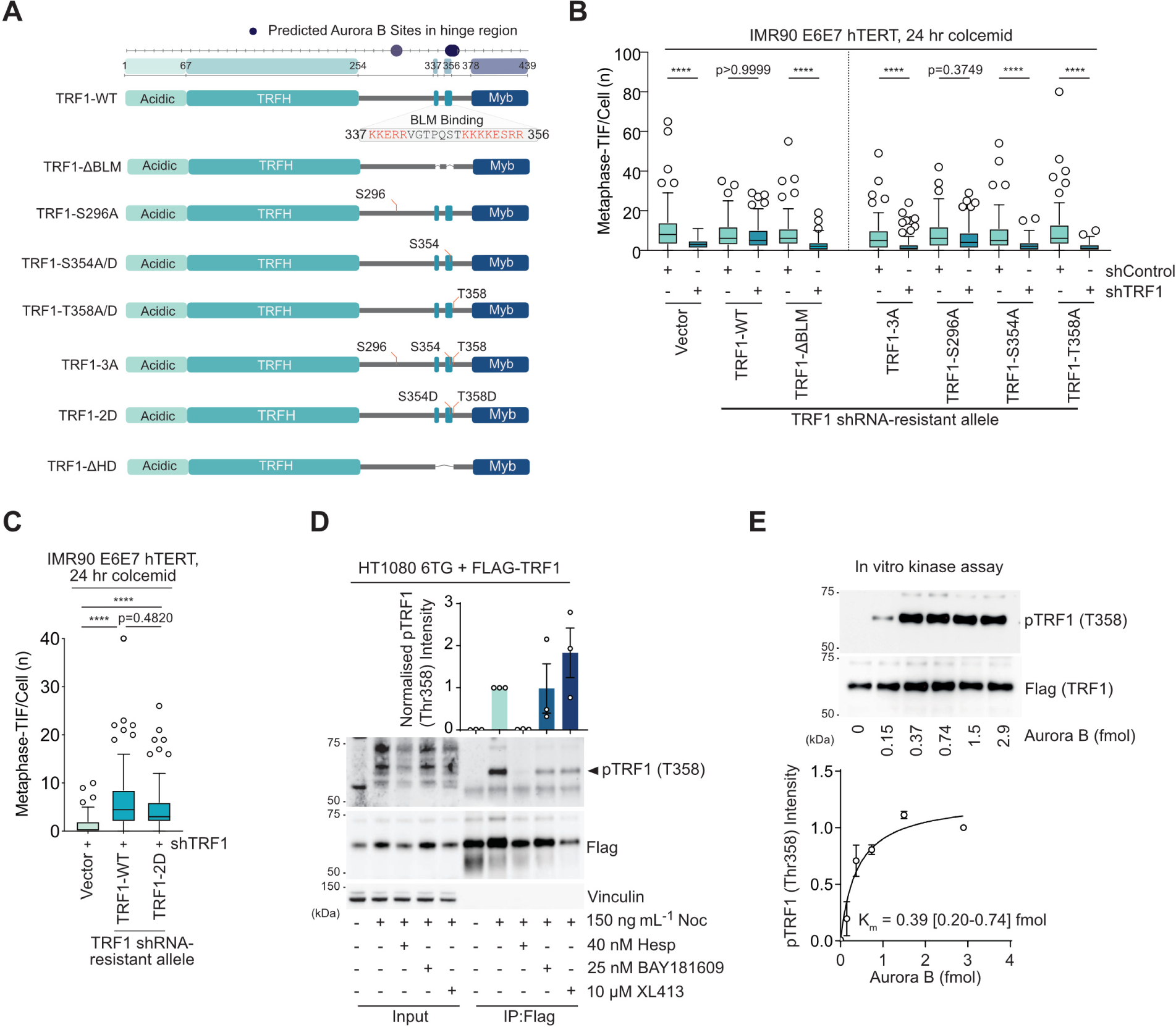
MAD-telomere deprotection requires TRF1 phosphorylation by AURKB. **A.** TRF1 alleles used in this study. Predicted AURKB sites in the hinge region and BLM binding domain are shown. The deleted amino acids within the FLAG-TRF1^ΔBLM^ variant are indicated in red. **B, C.** Metaphase-TIF following 24 hours of 100 ng mL^−1^ colecmid in IMR90 E6E7 hTERT fibroblasts expressing control or TRF1 shRNA and vector or shRNA-resistant TRF1 alleles (mean +/− s.e.m., n = 3 biological replicates of 30 metaphases per replicate, all data points combined into a Tukey boxplot, Kruskal-Wallis followed by Dunn’s multiple comparisons test, ****p<0.0001). **D.** Below, representative immunoblots of Flag immuno-precipitates from TRF1 shRNA HT1080 6TG cells expressing shRNA-resistant WT Flag-TRF1. Cells were synchronized with a thymidine block, released, and treated with 150 ng mL^−1^ nocodazole (Noc) where indicated, in the presence or absence of indicated kinase inhibitors for 16 hours. The pTRF1-T358 band is indicated with an arrow. Above, quantitation of normalized anti-TRF1-pT358 immunoblot signal (mean +/− s.e.m., n = 3 biological replicates). **E.** Above, example of an anti-TRF1-pT358 immunoblot measuring in vitro AURKB kinase assay on purified Flag-TRF1. Below, quantitation of anti-TRF1-pT358 intensity and the resulting Km (Graph is mean +/− s.e.m., n = 3 biological replicates, K_m_ is 0.39 (95% CI: 0.2-0.74) fmol).

We attempted to generate phospho-specific antibodies against these potential TRF1 phospho-sites. The phospho-TRF1-Ser354 (pTRF1-S354) antigen failed to produce specific antibody titres. Phospho-TRF1-Thr358 (pTRF1-T358), however, produced an antibody that reliably detected a band of approximately 60 kDa, corresponding to the size of Flag-TRF1 (Fig. S2F). The band was detected in anti-Flag immuno-precipitates from shTRF1 and Flag-TRF1^shR^ expressing HT1080 6TG cells enriched for mitotic arrest through G1/S synchrony and release into nocodazole for 16 hours (Fig. S2F). The band was absent from Flag-TRF1^shR^ immuno-precipitates from asynchronous cells, Flag-TRF1^shR^-T358A immuno-precipitates under all experimental conditions, and in alkaline phosphatase treated immuno-precipitates from Flag-TRF1^shR^ cells enriched for mitotic arrest (Fig. S2F). These data confirm antibody specificity and reveal accumulation of pTRF1-T358 during mitotic arrest.

To explore potential AURKB regulation of pTRF1-T358, HT1080 6TG cultures were G1/S synchronized, released, and enriched for mitotic arrest with 16 hours of 150 ng mL^−1^ Nocodazole in the presence or absence of mitotic kinase inhibitors (Fig 2D). The AURKB inhibitor Hesperadin was previously demonstrated to suppress MAD-TIF formation at 40 nM concentration^22^. Under these mitotic arrest conditions, 40 nM Hesperadin prevented pTRF1-T358 from exceeding interphase levels, while inhibitors against BUB1 (BAY1816032)^43^ or CDC7 (XL413)^44^ produced no effect (Fig. 2D). Additionally, low concentrations of AURKB vigorously phosphorylated purified Flag-TRF1 in vitro as measured by the anti-pTRF1-T358 antibody (Fig. 2E). Collectively the data indicate that TRF1 is required for MAD-telomere deprotection. Further, the data demonstrate that the role of TRF1 in this phenomenon is regulated through AURKB phosphorylation of TRF1 at T358 and potentially S354.

### Phosphorylated TRF1 recruits the CPC through Survivin

The TRF1 hinge region interacts with multiple proteins including BLM^40^. To identify proteins that interact with the AURKB modified TRF1 residues, we chemically synthesised phosphopeptides corresponding to pTRF1-S354 and pTRF1-T358. pTRF1-S296, shown above to not participate in MAD-telomere deprotection, and the known ATM site pTRF1-S367^45^ were included as controls. The phosphopeptides, and their respective non-phosphorylated controls, were immobilised on streptavidin beads and incubated with lysates from HT1080 6TG cultures enriched for mitotic arrest through G1/S synchrony and release for 16 hours in nocodazole (Fig. 3A-B). LC-MS/MS of proteins recovered by TRF1 peptide pulldown did not show enrichment of BLM nor its binding partners. Instead, we identified enrichment of the CPC component Survivin (BIRC5) in both the pTRF1-S354 and pTRF1-T358 peptide samples relative to non-phosphorylated controls (Fig. 3C and S3). Other CPC members Borealin (CDCA8) and AURKB were only weakly detected across all peptide pulldowns. Within the pTRF1-S354 sample we also recovered the 14-3-3 complex that is known to bind promiscuously to phosphorylated peptides^46^ (Fig. S3).

**Figure 3:**
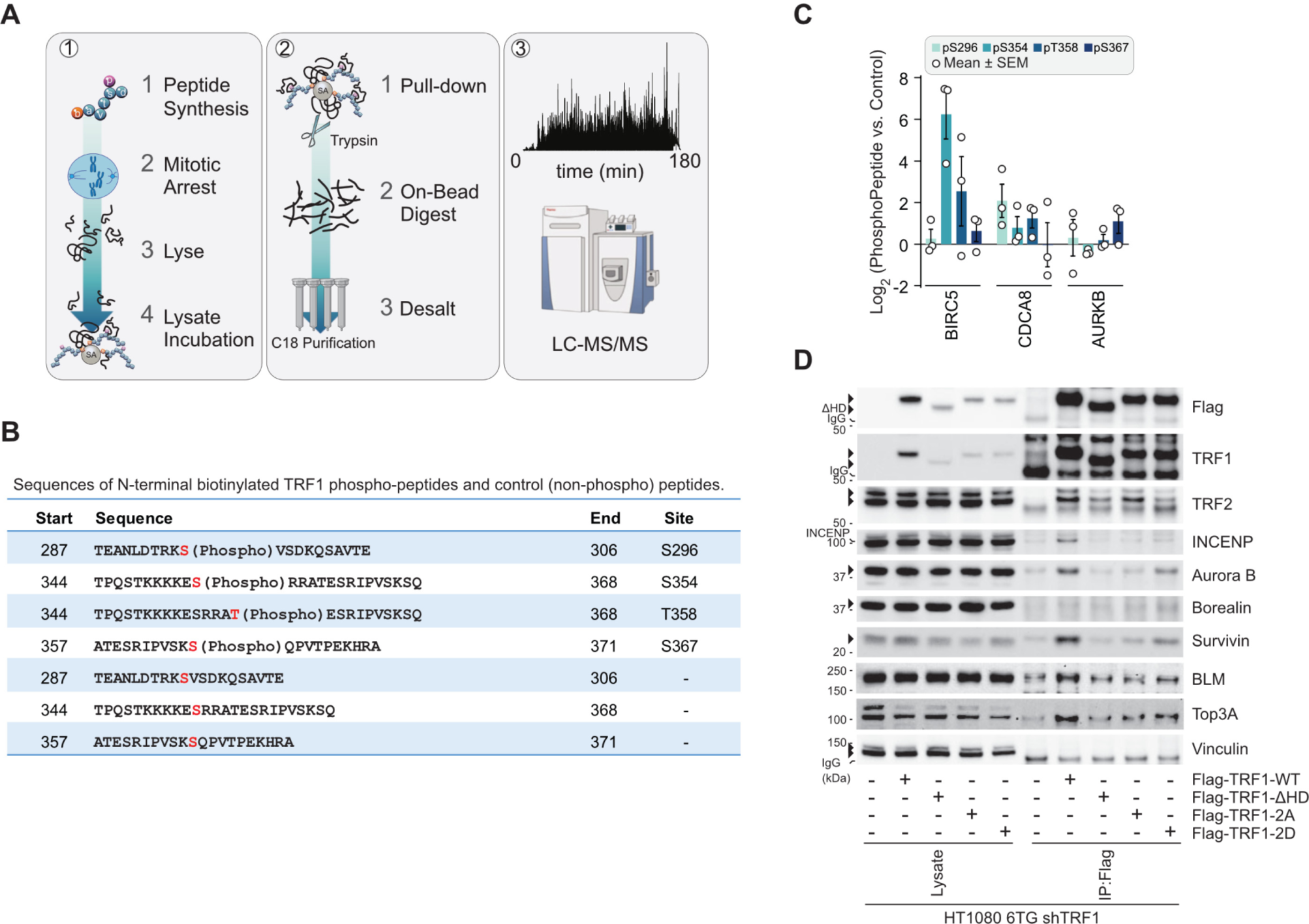
Survivin binds to TRF1 phosphorylated on Ser354 and Thr358. **A.** Schematic of TRF1 peptide pull-down and subsequent LC-MS/MS analysis for the experiment in Fig 3B, C and S3. To enrich for mitotic arrest, HT1080 6TG cells were synchronised with a thymidine block and released in the presence of 150 ng mL^−1^ Nocodalzole for 16 hours before sample collection and extract preparation. TRF1 peptides immobilized onto beads were incubated with HT1080 6TG mitotic extracts before LCMS/MS sample preparation and analysis. **B.** Summary of peptides used for pull-down analysis in Fig 3C and S3. Red indicates S296, S354, T358, and S367. **C.** Log_2_-fold change in Chromosome Passenger Complex proteins recovered from mitotically arrested HT1080 6TG extracts by TRF1 phosphopeptides (mean +/− s.e.m., n = 3 biological replicates). **D.** Immunoblots of anti-Flag immuno-precipitates from TRF1 shRNA HT1080 6TG cells expressing the indicated shRNA-resistant Flag-TRF1-WT or mutant alleles. Cells were synchronised with a thymidine block, released, and treated with 150 ng mL^−1^of nocodazole for 16 hours. TRF1-ΔHD is a deletion of residues 336-367 (representative of n = 3 biological replicates is shown).

To evaluate in a cellular context, we performed Flag co-IP in lysates from HT1080 6TG cells expressing TRF1^shR^-WT or mutant alleles (Fig 3D). Cells were enriched for mitotic arrest through G1/S synchrony and released into nocodazole for 16 hours before lysate collection. Interaction between TRF1 and the CPC components Survivin, INCENP, and AURKB were abrogated by deletion of the TRF1 hinge region (Flag-TRF1^shR^-ΔHD, aa336-367) that includes the S354 and T358 residues, and with double phospho-null Ala substitutions on S354 and T358 (Flag-TRF1^shR^-2A). CPC recovery was partially rescued by Flag-TRF1^shR^-2D carrying dual phospho-mimetic Asp substations at S354 and T358 (Flag-TRF1^shR^-2D)(Fig. 3D). Recovery of BLM and its binding partner TOP3A were also reduced in Flag precipitates from Flag-TRF1^shR^-ΔHD and Flag-TRF1^shR^-2A expressing cells and did not display obvious rescue in samples from Flag-TRF1^shR^-2D cells (Fig. 3D). pTRF1-S354 and pTRF1-T358 thus constitute a novel Survivin binding site. This region overlaps with the BLM-binding motif on TRF1, indicative of possible competition between BLM and CPC for TRF1 interaction. Survivin binding to AURKB-modified TRF1 suggests functional significance of CPC retention on Shelterin during mitotic arrest, potentially leading to the modification of other telomere proteins.

### AURKB phosphorylates the TRF2 basic domain to promote MAD telomere deprotection

MAD-TIF are ATM-dependent, suppressed by TRF2 over-expression, and enhanced by partial TRF2 depletion^25,26^; indicative of a central role for TRF2 in phenomenon regulation. We thus postulated AURKB may also modify TRF2. In silico analysis identified Ser62 and Ser65 in the human TRF2 basic domain as potential AURKB consensus sites (Fig. 4A). This was intriguing as the N-terminal TRF2 basic domain was previously suggested to protect t-loop junctions^32,33^.

**Figure 4.**
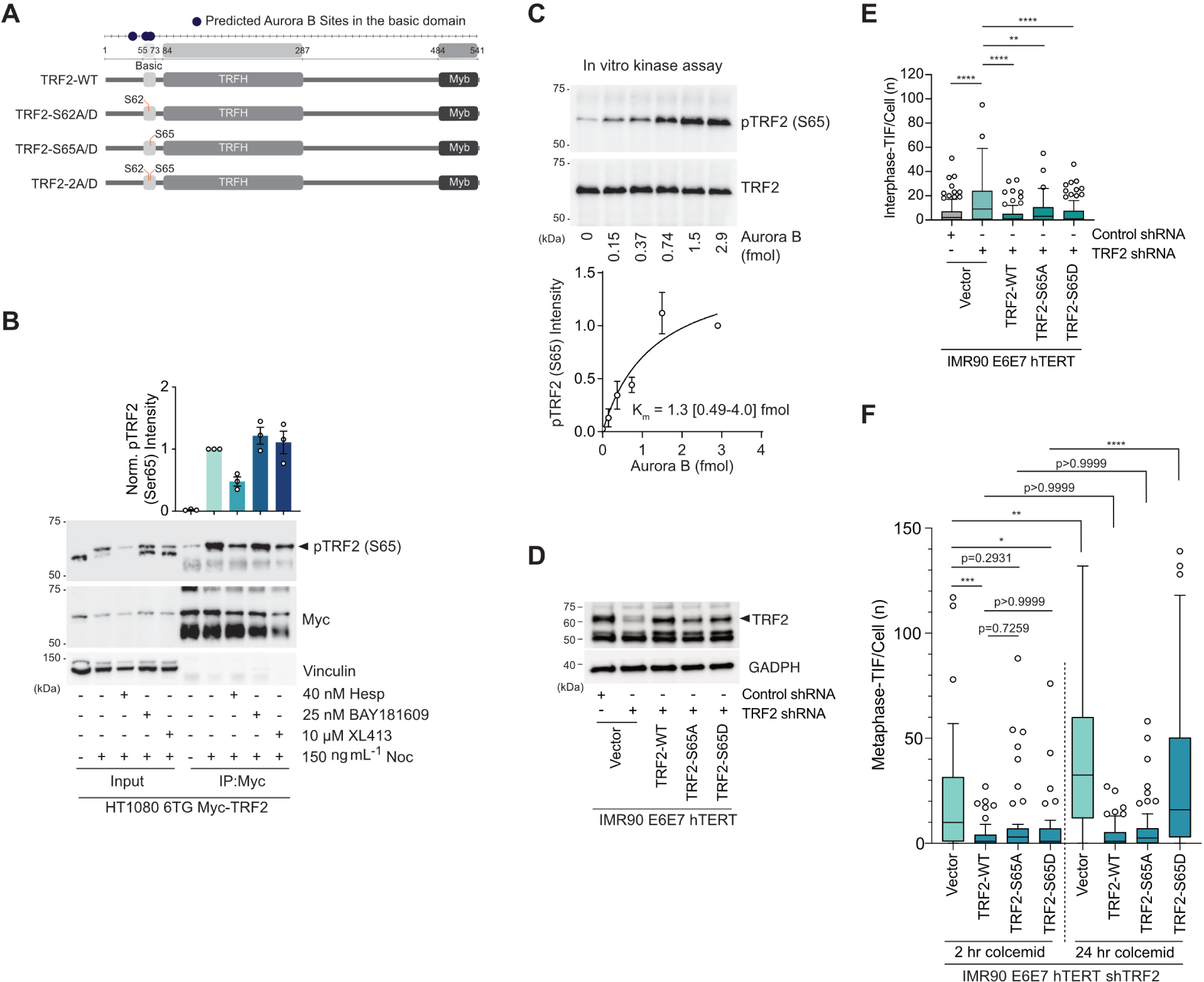
AURKB phosphorylates the TRF2 basic domain to promote MAD-telomere deprotection. **A.** TRF2 alleles used in this study. Predicated AURKB sites in the basic domain are indicated. **B.** Below, representative immunoblots of Myc immuno-precipitates from TRF2 shRNA HT1080 6TG cells expressing Myc-TRF2-WT. Cells were synchronized with thymidine block, released in the presence or absence of 150 ng mL^−1^ nocodazole (Noc) and the indicated kinase inhibitors for 16 hours. Above, quantitation of normalized anti-TRF2-pS65 immunoblot signal (mean +/− s.e.m., n = 3 biological replicates). **C.** Above, example of anti-TRF2-pS65 immunoblot measuring in vitro AURKB kinase assay on purified Myc-TRF2. Below, quantitation of anti-TRF2-pS65 intensity and the resulting Km (Graph is mean +/− s.e.m.., n = 3 biological replicates, Km is 1.3 (95% CI: 0.49-4.0) fmol). **D.** Representative immunoblots of whole cell extracts derived from IMR90 E6E7 hTERT cells transduced with control or TRF2 shRNAs and the indicated TRF2 wild type or mutant alleles. **E, F.** Quantitation of interphase-TIF (E) and metaphase-TIF (F) in TRF2 shRNA IMR90 E6E7 hTERT fibroblasts expressing TRF2-WT or the indicated TRF2 variants. For (E), n = 3 biological replicates of 45 nuclei per replicate, compiled into a Tukey boxplot, Kruskal-Wallis followed by Dunn’s multiple comparisons test, ****p<0.0001. For (F), cells were treated with 2 or 24 hours of 100 ng mL^−1^ colcemid (n = 3 biological replicates of 15 and 30 metaphases per replicate for 2 hours and 24 hours colcemid, respectively, compiled into a Tukey boxplot, Kruskal-Wallis followed by Dunn’s multiple comparisons test, ****p<0.0001).

We attempted to generate phospho-specific antibodies and failed for phospho-TRF2-Ser62 (pTRF2-S62). Anti-phospho-TRF2-Ser65 (pTRF2-S65), however, returned positive data. For these experiments we depleted endogenous TRF2 via shRNA and expressed exogenous TRF2-WT or mutant alleles. In whole cells extracts from HT1080 6TG cultures enriched for mitotic arrest with G1/S synchrony and released into nocodazole for 16 hours (Fig. S4A, Input), pTRF2-S65 revealed a band of approximately 60 kDa in Myc-TRF2-WT but not Myc-TRF2-S65A expressing cells. (Fig. S4A, Input). Immunoprecipitation of Myc-TRF2 from mitotically arrested cells revealed an enhanced band of the same molecular weight (Fig. S4A, IP:myc). We note a band of the same molecular weight, but less intense, was detectable following IP of Myc-TRF2-S65A from interphase and mitotically arrested cultures. We anticipate the pTRF2-S65 antibody may weakly recognize other phospho-sites on purified myc-TRF2, potentially Ser62, when Ser65 is no longer available for modification (Fig. S4A, IP:myc).

Consistent with pTRF2-S65 phosphorylation via AURKB, Hesperadin pre-treatment of cultures enriched for mitotic arrest reduced the intensity of the pTRF2-S65 band from Myc-TRF2 immunoprecipitates 52.1 ± 12.7% (mean ± s.d.) (Fig. 4B). TRF2 phosphorylation was unaffected by treatment with the Bub1 or CDC7 inhibitors BAY1816032 or XL413 (Fig. 4B). Additionally, in vitro incubation of immunoprecipitated Myc-TRF2 with recombinant AURKB produced substantial TRF2 phosphorylation as detected with the anti-pTRF2-S65 antibody, albeit with slower kinetics than TRF1 phosphorylation (Fig. 4C).

To explore the roles of TRF2-S62 and -S65 in mitotic telomere protection we shRNA depleted endogenous TRF2 in IMR90 E6E7 hTERT and complemented with ectopic wild-type or mutant TRF2 (Fig. 4D). Congruent with prior results^25,29^: TRF2 depletion conferred interphase-TIF and metaphase-TIF with 2 hours of 100 ng mL^−1^ colcemid^29^, consistent with passage of interphase-TIF into mitosis (Fig. 4E, F and S4B-E); TRF2 depletion also exacerbated the number of metaphase-TIF observed with 24 as compared to 2 hours of colcemid, indicative of an enhanced MAD-TIF response^29^ (Fig. 4F, S4E); and overexpression of ectopic TRF2-WT in a TRF2 shRNA background suppressed interphase-TIF and metaphase-TIF under all colcemid conditions^25,29^ (Fig. 4E, F and S4C-E). Notably, all TRF2-S62 and/or 65 mutant alleles rescued interphase protection in a TRF2 shRNA background (Fig. 4E and S4C, D, F)

Examination of TRF2-S65 revealed straightforward evidence consistent with phosphorylation of this residue being requisite for MAD-telomere deprotection. With two hours colcemid there was no difference in metaphase-TIF between WT-TRF2, TRF2-S65A and TRF2-S65D (Fig. 4F). Whereas the phospho-mimetic TRF2-S65D, but not phospho-null TRF2-S65A, was permissive to metaphase-TIF with 24 hours colcemid. Analysis of the TRF2-S62 mutants, however, revealed a contextually subtle phenotype. While all TRF2-S62 mutants, including TRF2-S62A/S65A (TRF2-2A) and TRF2-S62D/S65D (TRF2-2D), supressed interphase-TIF, all TRF2-S62 mutations also permitted metaphase-TIF with both two and 24 hours of colcemid (Fig. S4F, G). This indicated that within the context of TRF2-S62 mutation, metaphase-TIF observed with two hours of colcemid are not interphase-TIF passed into mitosis. Instead, the metaphase-TIF observed with two hours of colcemid in TRF2-S62 mutants represent MAD-TIF that arise through accelerated kinetics. We anticipate the sensitivity of TRF2-S62 to both phospho-null and -mimetic substitutions indicates the central importance of this residue in mitotic telomere protection. Collectively, the findings implicate the TRF2 basic domain in mitosis specific protection, and reveal this protective capacity is attenuated by AURKB-dependent modification.

### Attenuation of the TRF2 basic domain promotes MAD t-loop opening

To determine how TRF2 basic domain attenuation impacts mitotic t-loops, we sought to directly visualize telomere macromolecular structure through near super-resolution Airyscan microscopy^11^. For this purpose, we used *Trf2^Floxed/Floxed^ Rosa26-CreERT2 pBabeSV40LT* murine embryonic fibroblasts (hereafter abbreviated *Trf2^F/F^ CreER LgT* MEFs) which contain long telomeres that are easier to resolve with enhanced fluorescence microscopy^12^. *Trf2^F/F^ CreER LgT* MEFs were complimented with retroviral vectors expressing ectopic human TRF2 alleles and selected for transduction (Fig. S5A) Following endogenous *mTrf2* deletion with 4-OHT we assessed interphase-TIF and metaphase-TIF in cells arrested with Nocodazole for 14 hours. Consistent with IMR90 E6E7 hTERT, all hTRF2 mutant alleles restored interphase telomere protection in 4-OHT treated *Trf2^F/F^ CreER LgT* MEFs (Fig 5A and S5B). Also congruent between IMR90 E6E7 hTERT and 4-OHT treated *Trf2^F/F^ CreER LgT* MEFs, hTRF2-WT and hTRF2-S65A suppressed MAD-TIF while the hTRF2-2A and hTRF2-2D alleles harboring S62 mutations did not (Fig 5B and S5B). We note a subtle difference between MEFs and IMR90 E6E7 hTERT, as hTRF2-S65D was susceptible to MAD-TIF in the human but not the murine context (Fig 4F, 5B).

**Figure 5.**
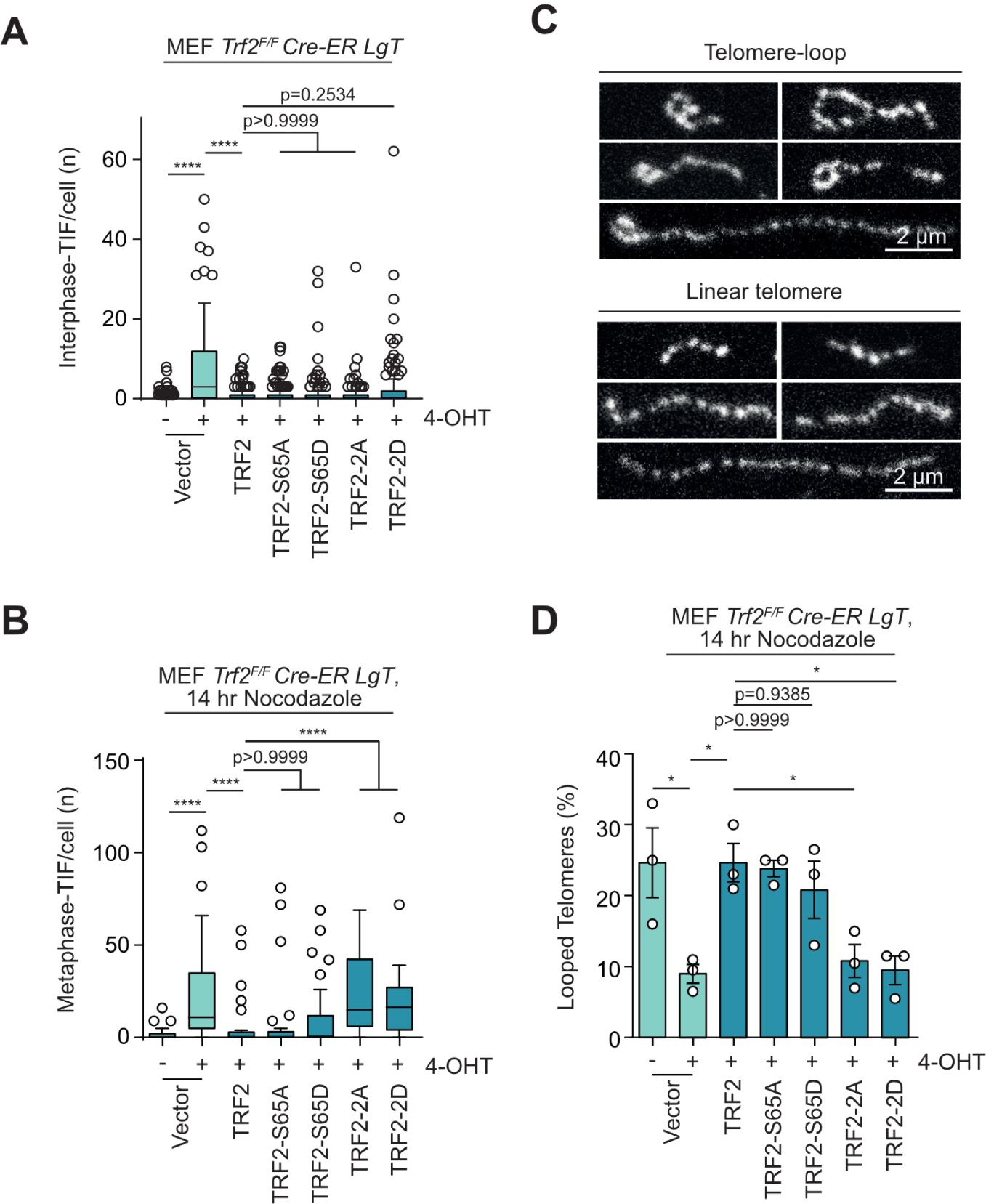
TRF2 basic domain modification during mitotic arrest promotes t-loop opening. **A, B.** Interphase-TIF (A) and Metaphase-TIF (B) in *Trf2^F/F^ Cre-ER LgT* MEFs expressing TRF2-WT or the indicated TRF2 variants. Where indicated endogenous TRF2 was deleted by 4-OHT addition 36 hours prior to sample fixation. In (B) cells were treated with 14 hours of 400 ng mL^−1^ Nocodazole prior to sample preparation (mean +/− s.e.m., n = 3 biological replicates of 90 cells per replicate, compiled into a Tukey box plot, Kruskal-Wallis followed by Dunn’s multiple comparisons test, ****p<0.0001) **C.** Representative examples of telomere macromolecular structure as visualized by AiryScan microscopy from *Trf2^F/F^ Cre-ER LgT* MEFs treated with 400 ng mL^−1^ Nocodazole for 14 hours and collected by mitotic shake-off. Samples were then trioxsalen cross-linked in situ, and the chromatin spread on coverslips through cytocentrifugation before telomere FISH labelling^11^. Scale bar, 2 µm. **D.** Quantification of looped telomeres from *Trf2^F/F^ Cre-ER LgT* MEFs expressing the indicated TRF2 alleles in mitotically arrested samples prepared and shown as in (C) and Fig S5C. Where indicated endogenous TRF2 was deleted by 4-OHT addition 36 hours prior to sample fixation (mean +/− s.e.m., n = 3 biological replicates of 200 telomeres per replicate, Ordinary one-way ANOVA followed by Šídák’s multiple comparisons test, F=6.463, DF=(6, 14)).

To assess mitotic telomere configuration, we 4-OHT treated *Trf2^F/F^ CreER LgT* MEFs and 36 hours later enriched cultures for mitotic arrest with 14 hr of 400 ng mL^−1^ nocodazole. Mitotic cells were collected via shake off and interstrand DNA crosslinks were introduced in situ with trioxsalen and UV light^6^. Interstrand crosslinking is required to maintain t-loop structure^6,11,12^. The chromatin was then cytocentrifuged onto glass coverslips in the presence of a mild detergent to decompact the telomeres prior to staining with telomere FISH^11,12^. Images were collected by airyscan microscopy and telomeres scored as looped or linear in blinded samples (Fig 5C and S5C).

Quantitation of telomere configuration in mitotically arrested cells revealed approximately 20% looped mitotic telomeres in *Trf2^F/F^ CreER LgT* MEFs with endogenous *Trf2*, and in 4-OHT treated samples expressing hTRF2-WT, hTRF2-S65A, or hTRF2-S65D (Fig 5D). Twenty-percent looped telomeres is consistent with prior observations of cells containing wholly protected telomeres^11,12,15,39^, and likely represents the limitation of trioxsalen crosslinking of t-loop junctions in situ^11^. Conversely, we found a diminished frequency of looped telomeres in 4-OHT treated *Trf2^F/F^ CreER LgT* MEFs transduced with vector, hTRF2-2A, or hTRF2-2D (Fig 5D). The inverse correlation observed between metaphase-TIF and t-loops during mitotic arrest (compare Fig 5B and D) is consistent with mitotic attenuation of the TRF2 basic domain promoting MAD-TIF through telomere linearization.

### BTR activity is required for MAD telomere deprotection

The above observations are consistent with the TRF2 basic domain counteracting BLM-dependent MAD-telomere deprotection. To determine which attributes of BLM promote MAD-TIF, we shRNA depleted BLM in IMR90 E6E7 hTERT and ectopically expressed shRNA resistant BLM alleles (BLM^shR^, Fig. 6A). In cultures treated for 24 hours with 100 ng mL^−1^ colcemid, BLM^shR^-WT promoted MAD-TIF as did the BLM^shR^-C1055S ATPase mutant that is mislocalized in interphase cells and defective for DNA duplex unwinding^47^ (Fig. 6A, B). Conversely, MAD-TIF were not rescued by mutations in the helicase (BLM^shR^-D795A)^48^ and HRDC domains (BLM^shR^-K1270V^49^) that attenuate dHJ dissolution (Fig. 6A, B).

**Figure 6.**
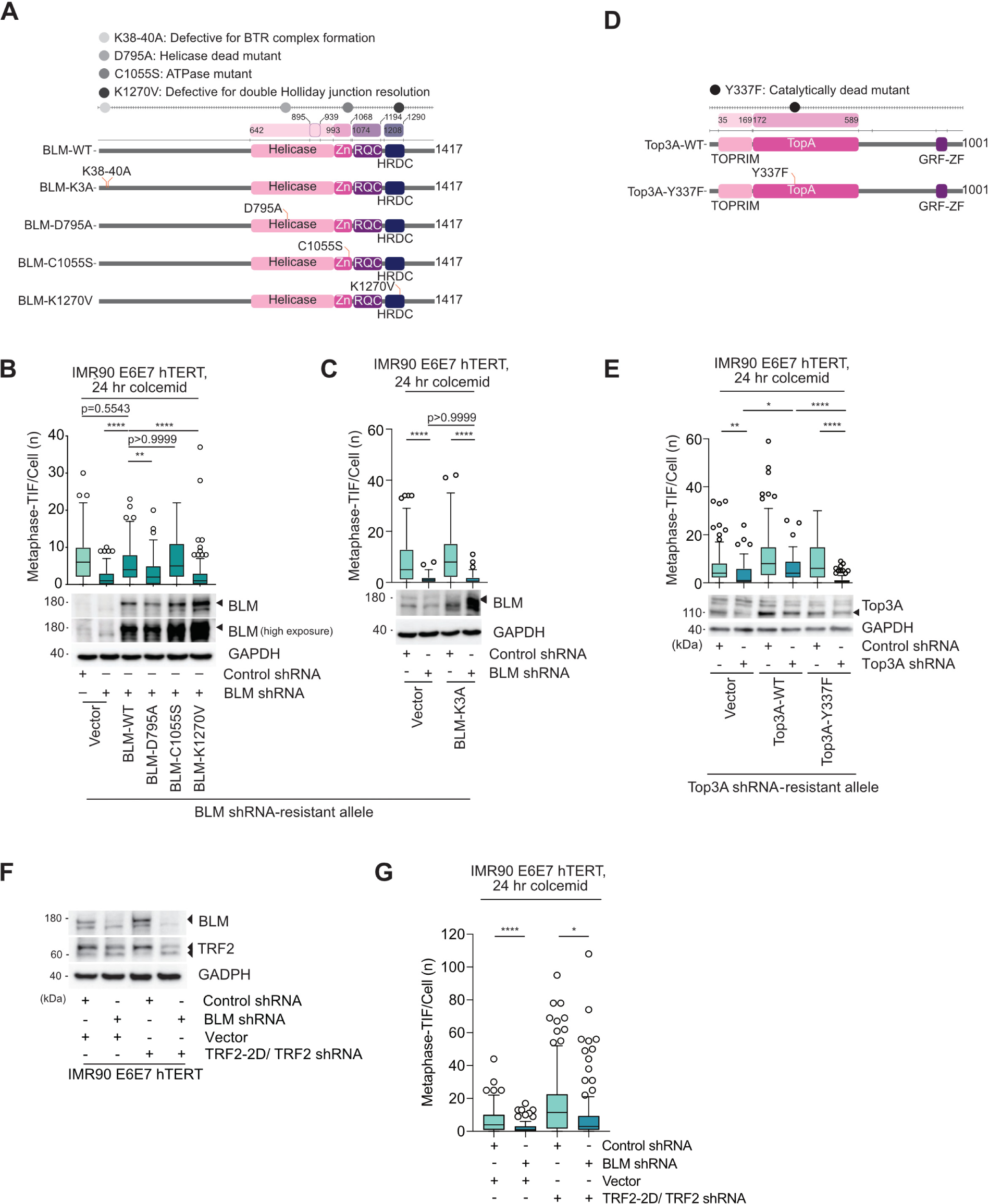
TRF2 phosphorylation promotes BTR-dependent MAD telomere deprotection. **A.** BLM alleles, and the respective compromised function of the indicated mutants, used in this study. **B, C.** Above: Metaphase-TIF following 24 hours of 100 ng mL^−1^ colcemid in IMR90 E6E7 hTERT fibroblasts expressing Control or BLM shRNA and shRNA-resistant BLM alleles (mean +/− s.e.m., n = 3 biological replicates of 30 metaphases per replicate, Kruskal-Wallis followed by Dunn’s multiple comparisons test, ****p<0.0001). Below: representative immunoblots of whole cell extracts derived from the cell cultures used in this experiment. **D.** Top3A alleles used in this study. **E.** Metaphase-TIF following 24 hours of 100 ng mL^−1^ colcemid in IMR90 E6E7 hTERT fibroblasts expressing Control or Top3A shRNA and vector or shRNA-resistant Top3A alleles (mean +/− s.e.m., n = 3 biological replicates of 30 metaphases per replicate, Kruskal-Wallis followed by Dunn’s multiple comparisons test, ****p<0.0001). Below: representative immunoblots of whole cell extracts derived from the cell cultures used in this experiment. **F.** Immunoblot of whole cell extracts derived from IMR90 E6E7 hTERT fibroblasts expressing Control, BLM or TRF2 shRNA, and vector or TRF2-2D (S62D and S65D). Cells were transduced five days prior to sample collection. Representative example of n = 3 biological replicates is shown. **G.** Metaphase-TIF following 2 or 24 hours of 100 ng mL^−1^ colcemid in IMR90 E6E7 hTERT fibroblasts expressing Control, BLM or TRF2 shRNA, and vector or TRF2-2D (mean +/− s.e.m., n = 3 biological replicates of 30 metaphases per replicate, Kruskal-Wallis followed by Dunn’s multiple comparisons test, ****p<0.0001).

Within the spectrum of BLM activities, dHJ dissolution is carried out by the BTR complex^36^. Expression of the BLM N-terminus Lys38-40 (BLM^shR^-K3A)^50^ mutant that disrupts interaction between BLM and other BTR components abolished MAD-TIF formation (Fig. 6C). As did individual depletion of the BTR components TOP3A, RMI1, or RMI2 (Fig. S6A, B). MAD-TIF were also rescued in TOP3A-depleted cells by TOP3A^shR^-WT but not the enzyme-dead mutant TOP3A^shR^-Y337F^51^ (Fig. 6D, E). As a control, we depleted FANCJ, a BLM associated helicase that operates outside of BTR^52^ (Fig. S6A). This had no effect on MAD-TIF (Fig. S6B). Together this causal evidence links BTR catalytic activity to MAD-telomere deprotection.

We predicted AURKB-modification of the TRF2 basic domain enabled BTR-dependent dissolution of t-loop junctions, thereby linearizing chromosome ends to promote telomere DDR activation. In agreement, BLM depletion suppressed MAD-TIF in TRF2 shRNA IMR90 E6E7 hTERT cultures that also expressed TRF2-2D (Fig. 6F, G). BTR-dependent dissolution disentangles dHJs without nucleolytic cleavage. Should t-loop junctions assume a dHJ configuration, we anticipate BTR-dependent dissolution will linearize the telomeres without the rapid telomere deletions that occur when t-loop junctions are resolved through nucleolytic pathways^32,33^. In support, we observed no reduction in the telomere lengths of DDR-positive telomeres that arise through MAD-telomere deprotection when compared to their protected sister chromatid (Fig S6C, D). AURKB therefore phosphorylates TRF2-S65, and potentially S62, during mitotic arrest to compromise telomere protection against BTR. We interpret the data to indicate BTR facilitates t-loop junction dissolution without telomere shortening to promote MAD-TIF.

### TRF2 modification impacts MAD lethality

A minority of cells experiencing mitotic arrest perish because of MAD telomere deprotection^22,24^. Altering TRF2 phospho-status should therefore affect a subset of mitotic death events. To test this directly, we depleted endogenous TRF2 in HT1080 6TG cultures and complemented the cells with TRF2-WT or TRF2 carrying Ser65 mutations (Fig. 7A). Under lethal replication stress, HT1080 6TG cells exhibit mitotic arrest and mitotic death signaled in part through MAD-telomere deprotection^24^. We treated the cultures with 1 µM of the DNA polymerase inhibitor aphidicolin and measured cell outcomes through widefield live imaging (Fig. 7B and S7A). Metaphase-TIF analysis was performed 40 hours after aphidicolin addition (Fig. 7B and S7B).

**Figure 7.**
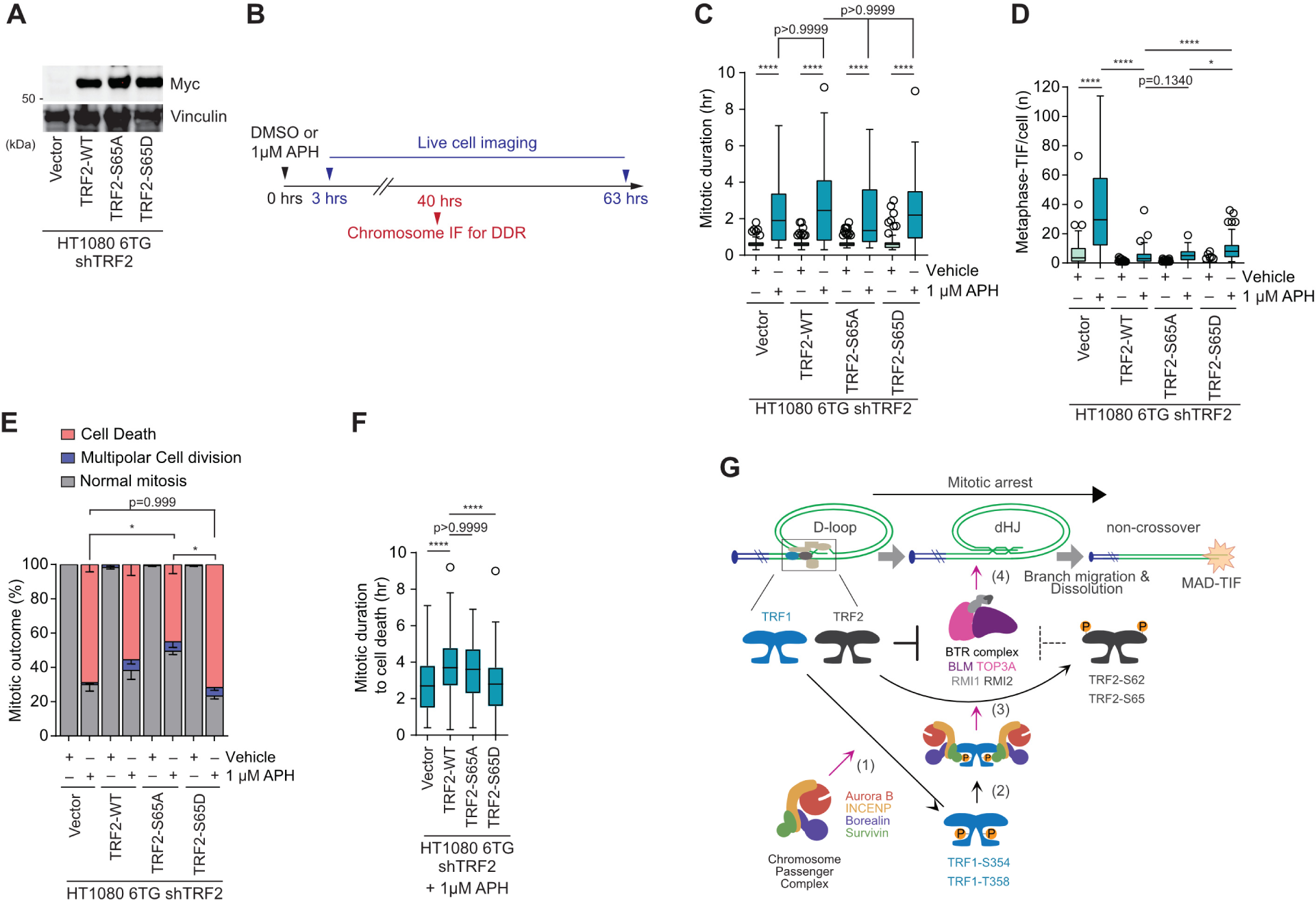
TRF2 phosphorylation promotes mitotic death during mitotic arrest. **A.** Immunoblots of whole cell extracts derived from HT1080 6TG shTRF2 cells expressing the indicated Myc-TRF2 alleles (representative example of n = 3 biological replicates). **B.** Timeline of the experimentation in panels C-F. **C.** Mitotic duration for TRF2 shRNA HT1080 6TG cells expressing the indicated Myc-TRF2 alleles. Cells were observed for 60 hours in cultures treated with or without 1µM Aphidicolin (APH) (mean +/− s.e.m., n = 3 biological replicates of 60 mitotic cells per replicate, Kruskal-Wallis followed by Dunn’s multiple comparisons test, ****p<0.0001). **D.** Quantitation of metaphase-TIF from TRF2 shRNA HT1080 6TG cultures expressing the indicated Myc-TRF2 alleles. Where indicated, cultures were treated with 1µM Aphidicolin (APH) for 40 hours before sample collection (n = 3 biological replicates of 30 metaphases per replicate compiled into a Tukey Box Plot, Kruskal-Wallis followed by Dunn’s multiple comparisons test, ****p<0.0001). **E.** Mitotic outcomes for TRF2 shRNA HT1080 6TG cells expressing the indicated Myc-TRF2 alleles. Cells were observed for 60 hours cultures treated with or without 1µM Aphidicolin (APH) (mean +/− s.e.m., n = 3 biological replicates of 60 mitotic cells per replicate, Ordinary one-way ANOVA followed by Šídák’s multiple comparisons test, F=6.931, DF=(3, 8), ****p<0.0001). **F.** Mitotic duration until death from the experiment in (E) (mean +/− s.e.m., n = 3 biological replicates of 60 mitotic cells per replicate, Kruskal-Wallis followed by Dunn’s multiple comparisons test, ****p<0.0001). **G.** Model of MAD telomere deprotection. (1) AURKB phosphorylates TRF1 at S354 and T358; (2) Survivn interacts with pS354 and pT358 to promote (3) AURKB phosphorylation of TRF2 at S62 and S65, enabling (4) BTR to dissolve t-loops.

Single cell analysis of the live imaging data identified that TRF2 expression status did not affect mitotic arrest duration induced by a lethal aphidicolin treatment (Fig. 7C). As reported previously, the enhanced metaphase-TIF in TRF2 shRNA cells treated with lethal replication stress were suppressed by ectopic TRF2-WT (Fig. 7D). Within aphidicolin treated cultures, cells expressing TRF2-S65D displayed significantly more metaphase-TIF than cultures expressing TRF2-WT or TRF2-S65A (Fig. 7D). We note that with aphidicolin metaphase-TIF can result from both interphase telomere replication stress carried into mitosis and MAD-telomere deprotection. Because aphidicolin induces a less severe mitotic arrest phenotype than microtubule poisons, we anticipate metaphase-TIF in aphidicolin-treated shTRF2 vector cells primarily result from interphase replication stress with a minor MAD-telomere deprotection component. The live imaging data revealed that mitotic cell death was attenuated by TRF2-S65A but not TRF2-S65D (Fig. 7E). Consistent with MAD-TIF accumulation signaling mitotic lethality^22,24^, the duration of mitotic arrest to mitotic death was elongated when MAD-telomere deprotection was suppressed by TRF2-S65A, but was not affected by TRF2-S65D (Fig. 7F). Together the data are consistent with AURKB-dependent TRF2 phosphorylation promoting the minor proportion of mitotic death events signaled through MAD telomere deprotection.

## DISCUSSION

Here we describe a mechanism of MAD telomere deprotection regulated by the CPC, shelterin, and BTR. Our data reveal that the CPC component AURKB phosphorylates TRF1 at Thr358 and potentially Ser354 specifically during mitotic arrest (Fig. 7G [1]). These phosphorylation events promote binding of TRF1 by the CPC through its Survivin subunit (Fig. 7G [2]). During mitotic arrest, AURKB also phosphorylates the TRF2 basic domain on Ser65 and potentially Ser62 (Fig. 7G [3]). AURKB-dependent attenuation of the TRF2 basic domain enables BTR dissolution of t-loop junctions, resulting in linear telomeres and a localized ATM-dependent DDR (Fig. 7G [4]), which contributes to mitotic death. Through identification of TRF1-dependent ATM regulation at somatic telomeres, and participation of TRF1 and TRF2 in an active process of telomere deprotection, we reveal an unappreciated complexity in shelterin function.

### MAD-telomere deprotection kinetics

MAD-telomere deprotection requires four or more hours of mitotic arrest^25^ and, as demonstrated here, involves AURKB-dependent modification of multiple shelterin factors. During normal cell division the CPC is subject to tight spatiotemporal regulation^35^. Mitotic arrest occurs during prometaphase or metaphase when the CPC is localized at kinetochores to regulate microtubule attachment and the spindle assembly checkpoint. We were unable to identify where shelterin and the CPC first interact. However, observation of multiple kinetochore factors in our APEX2-Flag-TRF1 interactomics dataset suggests TRF1 localization at kinetochores during mitotic arrest. In support, TRF1 was previously implicated in centromere function^53^. Further, respective mitotic TRF1 phosphorylation on Thr371 in human and Ser404 in mice by CDK1 and AURKB were proposed to dissociate TRF1 from the telomere substrate^54,55^. This may increase the pool of diffusive TRF1 in mitotic cells, facilitating stochastic and transient kinetochore interactions.

It is tempting to speculate that diffusive TRF1 stochastically interacts with kinetochores leading to CPC-directed phosphorylation of TRF1-S354 and/or T358. Phosphorylated TRF1, alone or in complex with the CPC via interaction through Survivin, may continue diffusive movement within mitotic cells. Eventually, the CPC-TRF1 complexes can interact with a telomere and promote further AURKB-dependent shelterin phosphorylation. Alternatively, or in parallel, diffusive CPC may instead stochastically interact with chromosome ends and phosphorylate telomere bound shelterin. The slow kinetics of MAD-telomere deprotection likely reflects the time required for such undirected movements to occur.

We were unable to conclusively determine the exact order of phosphorylation events occurring during MAD-telomere deprotection. However, AURKB phosphorylated TRF1-T358 with faster kinetics than TRF2-S65 in vitro, suggesting TRF2 phosphorylation is a rate limiting step. Additionally, mutational analysis was consistent with TRF2 modification being the terminal phosphorylation event. TRF2-S65A inhibited both interphase-TIF and MAD-TIF, whereas TRF2-S65D inhibited interphase-TIF but was permissive to MAD-TIF with typical kinetics. All TRF2-S62 mutants, however, suppressed interphase TIF, but were permissive to metaphase-TIF with two hours colcemid. This is consistent with accelerated MAD-TIF kinetics following TRF2-S62 alteration. TRF2-S62 is therefore crucial for basic domain protection and exquisitely sensitive to modification. Further, these findings demonstrate that the physiological role of the TRF2 basic domain is the provision of mitosis-specific protection.

We interpret the data to indicate that MAD-TIF are potentiated primarily through CPC recruitment to shelterin via TRF1, which enables subsequent TRF2 phosphorylation. Directly attenuating basic domain protection via TRF2-S62 mutation circumvents the need for upstream TRF1 recruitment and accelerates MAD-TIF kinetics. This also indirectly supports AURKB modification of its TRF2-S62 consensus sequence being requisite for MAD-telomere deprotection. Because TRF2-S65D fails to accelerate MAD-TIF kinetics, this argues that additional TRF1-directed and AURKB-dependent TRF2-S62 phosphorylation is required for MAD-TIF to form. Additionally, while the CPC and BLM binding motifs on TRF1 overlap, the phospho-mimetic TRF1-2D mutant rescues CPC but not BLM recovery in Co-IPs. This argues that MAD-telomere deprotection relies more on CPC and TRF1 interaction than BLM and TRF1 interaction. We anticipate the slow kinetics of a sequential TRF1, CPC, TRF2, BTR pathway results in continued telomere protection during normal mitoses.

### The TRF2 basic domain regulates mitotic telomere protection

Data from several publications support protection of t-loop junctions by the TRF2 basic domain. Biochemical evidence demonstrates that the TRF2 basic domain binds 3- and 4-way DNA junctions^31,32,34^, and also remodels oligonucleotides containing telomeric DNA into 4-way HJs^34^. Expression of a TRF2 mutant allele lacking the basic domain (TRF2^ΔB^) promotes rapid telomere deletions and free telomere DNA circles dependent upon homologous recombination (HR)^33^. Swapping the TRF2 basic domain with prokaryotic factors that bind 4-way DNA junctions restored TRF2 protection against such rapid telomere deletions^32^. Collectively the data indicate that when the TRF2 basic domain is absent, t-loop junctions are exposed to nucleolytic cleavage – a process termed t-loop HR^33^. T-loop HR is more pronounced in TRF2^ΔB^ cells with concurrent BLM knockout, and unchecked t-loop HR can shorten telomeres sufficiently to promote chromosome fusions and genome instability^32^.

Expression of exogenous TRF2^ΔB^, however, does not induce a strong interphase-TIF phenotype^33^. In agreement, all TRF2 basic-domain mutants examined here retained interphase protection, including the TRF2-S62 mutants that accelerated MAD-TIF formation. This argues that the physiological role of the basic domain is mitosis-specific protection. Understanding HR is informative to why mitosis-specific t-loop junction protection is required. During HR, strand-invasion can be followed by second-end DNA capture and dHJ formation^56^. HJs are resolved in mammalian cells through nucleolytic cleavage by the SLX4-SLX1-MUS81-EME1 complex^57^ or GEN1^58^; enzymes whose HJ resolution activity is spatiotemporally restricted to mitosis^59^. Alternatively, dHJs can be dissolved without cleavage through the BTR complex^36^ and BLM hyper-phosphorylation at the G2/M transition is required for full dissolution activity^60^. We anticipate that t-loop HR in mammalian interphase nuclei is tempered by inhibition or exclusion of HJ resolvase activity. When TRF2^ΔB^ cells enter mitosis, however, the exposed t-loop junctions are likely subject to nucleolytic resolution and/or BTR dissolution.

We did not observe quantitative telomere shortening associated with MAD-telomere deprotection. We also did not observe unequal sister telomere intensity in mitotic chromosome spreads, the traditional qualitative evidence of rapid telomere deletions^25,33,61^. T-loop HR therefore remains suppressed following AURKB modification of the TRF2 basic domain. We interpret our findings to indicate that the basic domain protects t-loop junctions from both BTR-dependent dissolution and nucleolytic cleavage resulting in t-loop HR. TRF2 modifications during mitotic arrest derepresses protection against BTR while maintaining protection against nucleolytic resolution. This enables t-loop junction dissolution, chromosome end linearity, and DDR activation without the risk of rapid telomere deletions.

### The molecular identity of t-loop junctions

The involvement of BTR in MAD-TIF formation is consistent with t-loop junctions assuming a dHJ configuration during mitotic arrest. It remains unclear, however, if dHJs are ubiquitous at the t-loop insertion point, or if these double four-way structures are only formed during mitotic arrest. We currently favor the hypothesis that t-loop junctions typically assume a 3-way D-loop or 4-way single HJ that is bound and stabilized by the TRF2 basic domain. AURKB modification likely disrupts electrostatic interactions between the basic domain and its structured DNA substrate, allowing for branch migration, dHJ formation, and dissolution via BTR. While a dHJ would provide greater interaction between the distal invading and proximal receiving telomere portions, dHJs would also require coordinated resolution ahead of oncoming replication forks in S-phase. The temporal dissolution of mitotic t-loops described here is reminiscent of RTEL1-dependent t-loop unwinding during S-phase^39^. Both pathways are restricted to a specific cell cycle window, executed by a helicase, and regulated through TRF2 phosphorylation. Notably, RTEL1 suppresses cross-over repair during HR through a proposed anti-recombinase mechanism that limits second end capture and dHJ formation^62^. Inhibiting telomeric RTEL1 localization in S-phase promotes t-loop HR^39,63^. This is consistent with negative outcomes occurring when dHJs are present at the t-loop insertion point. Future structural studies of shelterin bound to structured DNA will ultimately clarify t-loop junction identity.

### Shelterin-mediated telomere deprotection

Shelterin is typically conceptualized as a chromosome end protective complex where individual components inhibit specific DDR and/or double strand break repair pathways. TRF2 is attributed as the sole factor in somatic cells responsible for t-loop formation and ATM activation^15,16,30^. To our surprise, we found TRF1 is required for MAD-telomere deprotection. This revealed that shelterin is necessary for active telomere deprotection during mitotic arrest, and implicated TRF1 in somatic ATM and t-loop regulation under specific physiological conditions. The wider implication of our findings is that shelterin protective activities are more entwined than previously understood. Crosstalk between shelterin subunits may influence DDR outcomes previously considered under the control of a single telomere factor. Understanding crosstalk between shelterin components may inform future studies on aging-dependent telomere deprotection.

Finally, our study reveals shelterin is a dynamic complex that mediates both telomere protection and deprotection. Because NHEJ is repressed in mitosis^64^, and TRF2 remains bound to deprotected mitotic telomeres to directly inhibit end-joining upon mitotic exit^29^, there is no risk of telomere fusions from MAD-telomere deprotection^25^. MAD-telomere deprotection is therefore a highly coordinated mechanism that facilitates chromosome end linearity and telomere DDR activation without compromising genome stability through telomere shortening or fusion. Because mitotic arrest is a common response to genomic damage, including tumor-relevant replication stress and chemotherapeutic intervention^23,24^, we anticipate MAD-telomere deprotection evolved as a failsafe to promote mitotic death or p53-dependent G1 arrest in genomically damaged cells.

## Acknowledgements

We thank Will Hughes, Scott L. Page, and the CMRI ACRF Telomere Analysis Centre supported by the Australian Cancer Research Foundation for microscopy infrastructure and support; the Westmead Institute of Medical Research Flow Cytometry Centre, supported by the Cancer Council NSW and the Australian NHMRC for cell sorting; Mark Graham and George Craft and the CMRI Biomedical Proteomics Centre for Mass spectrometry infrastructure and support; the CMRI Peptide Synthesis Facility for their assistance; Andrea Ruelas-Gonzalez and Yuya Nishida for experimental support with metaphase-TIF analysis; Yumi Hayashi for assistance with molecular cloning; Ylli Doksani for critical reading of the manuscript; and Jan Karlseder, Didier Trono, and Robert A. Weinberg for sharing plasmids. Members of the Hayashi and Cesare labs are thanked for their suggestions and discussion. A.J.C. is supported by an Australian Research Council Future Fellowship (FT210100858). This project was supported by grants from the Australian NHMRC to A.J.C. and M.T.H (1106241) and A.J.C. (1162886); the Australian Research Council to A.J.C. (DP210103885, DP240101869); an Australian Medical Research Future Fund award to A.J.C. (2007488); Japan Foundation for Applied Enzymology to M.T.H.; Grant-in-Aid for Young Scientists (A) (16H06176) to M.T.H.; Grant-in-Aid for Scientific Research (B) (20H03183) to M.T.H.; and the Hakubi project grant to M.T.H.; and institutional funding from CMRI (to N.L. and A.J.C.) and Kyoto University (to F.I. and M.T.H.).

## Author contributions

D.R.Z., S.R., M.T.H, and A.J.C. conceived the study. D.R.Z. and S.R. performed most of the experiments and data analysis in this manuscript. S.R. completed the mass spectrometry and analysis and R.R.J.L. the t-loop imaging and analysis. A.B.R. performed the peptide synthesis. M.T.H and A.J.C. assisted with data analysis throughout. S.G.P and B.L. created and verified plasmids and cell lines and assisted with minor experimentation throughout. N.L., F.I., M.T.H., and A.J.C. supervised junior members of the research team and secured funding. S.R., D.R.Z., M.T.H., and A.J.C. wrote the manuscript with editorial input from R.R.J.L., S.G.P., B.L., N.L.,

## Declaration of interests

The authors declare no competing interests.

## METHODS

### Cell culture and treatment

IMR90 fibroblasts were provided by Fuyuki Ishikawa (Kyoto University), HeLa cells by Megan Chircop (CMRI), HT1080 6TG cells by Eric Stanbridge (University of California, Irvine), and *TRF2^Floxed/Floxed^ Rosa26-CreERT2 pBabeSV40LT* MEFs by Eros Lazzerini Denchi (National Cancer Institute)^13^. IMR90 E6E7 hTERT fibroblasts produced as described previously^65^. IMR90 E6E7 hTERT fibroblasts were grown in Dulbecco’s Modified Eagle Medium (DMEM) supplemented with 10% fetal bovine serum (S1810-500, biowest), 200 mM L-glutamine, 7.5% NaHCO_3_, 100 U mL^−1^ penicillin, streptomycin, and 5 μg mL^−1^ Plasmocin (InvivoGen) and maintained at 37 °C in 5% CO_2_ and 3% O_2_. HT1080 6TG, HeLa-APEX2, HeLa-APEX2-TRF1, and MEF cultures were grown in Dulbecco’s Modified Eagle Medium (DMEM) supplemented with 10% fetal calf serum (F9423, Sigma), GlutaMax and MEM non-essential amino acid solution (Gibco) and maintained at 37 °C in 7.5% CO_2_ and 3% O_2_. Cell Bank Australia verified cell line identity using short-tandem-repeat profiling, and cells were identified to be mycoplasma negative (MycoAlert, LT07-118, Lonza).

HT1080-6TG, HeLa-APEX2, HeLa-APEX2-TRF1, and MEF cells were synchronized in mitosis by treating cultures at 70% confluence with 2 mM Thymidine (Sigma) for 24 hours. Thymidine was washed out with warm PBS, replacing with fresh media containing 150 ng mL^−1^ Nocodazole (M1404, Sigma) for 14-16 hours to arrest cells in mitosis. Rounded mitotic cells were detached from culture dishes and interphase cells by mitotic shake-off. Where indicated, cells were treated with 100 ng mL^−1^ Colcemid (15212012, Gibco), 150 ng mL^−1^ Nocodazole (M1404, Sigma), 40 nM Hesperadin (S1529, Selleck Chemicals), 10 µM XL413 (24906, Cayman Chemical), or 25 nM BAY181609 (A19868, Cayman Chemical).

### Plasmid cloning

Plasmids used in this study are listed in Table 1. Complementary DNA encoding human TRF1 (NP_059523.2), the functional short isoform of TRF2 (XP_005256180.1, missing the first 42 a.a.)^66^, and BLM (NP_000048.1) were generously provided by Jan Karlseder. The nuclear short isoform of Top3A (Q13472-2) was artificially synthesized (Integrated NDA Technologies). All cDNAs were in-frame cloned downstream of blasticidin S-resistance gene and self-cleaving p2a sequences in a 3^rd^ generation lentiviral plasmid vector. Flag and myc tags were added during PCR. Mutations and deletions were introduced by site-directed PCR mutagenesis (TRF1, TRF2, BLM) or during artificial DNA synthesis (Top3A) and cloned by the recombination-based HiFi assembly strategy (NEB). Amino acid positions in each protein correspond to the indicated reference sequences.

APEX2 and APEX2-TRF1 vectors were created by assembling fragments into the pRRL-sin-cPPT-hPGK-eGFP-WPRE backbone (a kind gift from Leszek Lisowski). All constructs were cloned using PCR amplification of complimentary fragments and In-Fusion HD assembly (Takara). pRRL-hPGK-mCherry-P2A-PuroR-T2A-FLAG-APEX2 was created by PCR amplifying hPGK-mCherry-P2A-PuroR-T2A from a synthesised gene fragment purchased from GeneWiz and APEX2 from pcDNA3-FLAG-APEX2-NES (a gift from Alice Ting^67^, Addgene plasmid # 49386) followed by insertion into PCR linearised pRRL backbone. pRRL-hPGK-mCherry-P2A-PuroR-T2A-FLAG-APEX2-TRF1 was created following the same approach, by fusing the same fragments with TRF1 PCR amplified from pLPC-myc-His-BirA-human TRF1 (a kind gift from Roderick O’Sullivan^68^).

Short hairpin RNA against TRF1, TRF2, BLM, Top3A, RMI1, RMI2, and FANCJ were cloned into pLKO.1 vector by conventional restriction enzyme cloning of annealed oligo nucleotides. The shRNA target sequences were as follows: Control 5’-CCTAAGGTTAAGTCGCCCTCGCTC-3’, BLM 5’-TGCCAATGACCAGGCGATC-3’^69^, TRF1 5’-CCCAGCAACAAGACCTTAATA-3’^70^, TRF2 5’-GCGCATGACAATAAGCAGATT-3’^29^, Top3A 5’-GCTTCTCGAAAGTTGAGAATA-3’^71^, RMI1 5’-CGATCGAAGTATAGAGAGATT-3’ (TRCN0000158474), RMI2 5’-CCATGAAAGTATGTGGGAACT-3’ (TRCN0000143418), and FANCJ 5’-CGTCAGAACTTGGTGTTACAT3’ (TRCN0000049914). shRNA-resistant silent mutations were as follows: BLM 5’-c GCt AAc GAt CAa GCc ATt-3’, TRF1 5’-Ct CAa CAg CAg GAt tTg AAc A-3’, Top3A 5’-GCc agc aGg AAa cTc cGg ATc-3’, where lower the case denotes silent mutations. Exogenous TRF2 alleles did not carry shRNA-resistant silent mutations as exogenous TRF2 expression in the presence of TRF2 shRNA produces sufficient protein to rescue wild-type TRF2 function^29^. All plasmid sequences were confirmed by Sanger sequencing and are available upon reasonable request.

### Site-directed mutagenesis

Site directed mutagenesis of plasmids listed in Table 1 was carried out using KAPA HiFi HotStart premix (Roche) or Tks Gflex DNA polymerase (Takara Bio) by gradient touch-down PCR with cycler settings optimised for long-range synthesis. Mutagenesis primers are listed in Table 2 and manufactured by IDT or Eurofins genomics. Following successful amplification, and confirmation by agarose gel electrophoresis, mutagenised amplicons were digested with DpnI (New England Biolabs (NEB)) to remove template DNA, phosphorylated by T4 polynucleotide kinase (NEB), ligated with Blunt/TA master mix (NEB) or DNA ligation kit (Takara) and transformed into Stellar^TM^ competent cells (Takara Bio) or Mix & Go Stbl3 competent cells (Zymo Research). Following standard plasmid DNA preparation, mutagenesis was confirmed by Sanger sequencing.

### Viral packaging and transduction

Lentivirus particles for transduction of IMR90 E6E7 hTERT cells were produced by transfection of an expression plasmid with packaging and envelope plasmids gifted from Didier Trono (Addgene plasmid #12260) and Robert A. Weinberg (Addgene plasmid #8454), respectively, into HEK293FT or its derivative Lenti-X 293T cells (632180, Takara). Media was replaced after 24 hours and the viral supernatant was collected through filtration (0.45 µm pore, 25 mm, technolabsc inc.) at 48- and 72-hours post-transfection and stored at –80 °C until transduction. The viral supernatant was complemented with 8 μg mL^−1^ polybrene for transduction. Selection was carried out by using 10 μg mL^−1^ Blasticidin-S (Funakoshi) for at least five days to obtain cells stably expressing target proteins. For knockdown experiments, cells were transduced with lentivirus carrying shRNA target sequences for two days and selected with 1 μg mL^−1^ Puromycin (ChemCruz) for more than three days before experimental procedures.

For transduction of HeLa or HT080 6TG cells with APEX2, APEX2-TRF1, or shRNA vectors, high titre, purified pRRL/pLenti-derived lentiviral vectors were created by the CMRI Vector and Genome Engineering Facility. HeLa cultures were transduced with concentrated lentivirus at an MOI of 10 for 48 hours in 4 µg mL^−1^ polybrene (Sigma), then selected in normal growth media supplemented with appropriate antibody selection; APEX2 vectors 0.25 µg mL^−1^ puromycin, shRNA vectors 1 µg mL or 5 µg mL^−1^ Blasticidin-S or 400 µg mL^−1^ Hygromycin. Following expansion, APEX2 cells were sorted for high mCherry expression by the Westmead Institute for Medical Research Cytometry Facility (Sydney, Australia). Positively transduced cells were expanded and frozen in FBS with 10% DMSO at low passage. For FLAG-TRF1, high titre lentiviral vectors created by the CMRI Vector and Genome Engineering Facility were added to HT1080-6TG cell cultures supplemented with 4 µg mL^−1^ polybrene for 48 hours, then selected in normal growth media supplemented with 1 µg mL^−1^ puromycin for 72 hours.

Retroviral particles were produced by transfecting pLPC plasmids into low passage Phoenix cells (ATCC) at 90% confluence using Lipofectamine3000 (ThermoFisher) as per the manufacturer’s instructions. Viral supernatants were removed at 24- and 48-hours post-transfection, filtered through 0.45 µm syringe filters and added to target cells in media containing 4 µg mL^−1^ polybrene. Cells were grown for 48 hours following retroviral transduction before selecting with 1 µg mL^−1^ of Puromycin (ChemCruz) for more than 72 hours before experimentation. All retro/lentiviral transduced cells were maintained for short-term culture in 50% concentration of selection antibiotic to retain transgene expression.

### APEX2-TRF1 proximity biotin-labelling

We added 500 µM Biotin-Phenol (Iris Biotech, LS-3500) in a suspension of culture media containing 150 ng mL^−1^ Nocodazole ± 40 nM Hesperadin (mitotic cells only) for 30 min. Cells were resuspended in 5 mL of media and 1 mM hydrogen peroxide was added to the cells for 60 sec to initiate APEX2 labelling. Reactions were quenched with 3x 3 mins washes of quench solution containing 20 mM Sodium Ascorbate (Sigma), 10 mM Trolox (Sigma) and 20 mM Sodium Azide (Sigma). The cells were washed with PBS and frozen at –80 °C as a dry pellet, prior to streptavidin pull-down. For attached G1/S cells, all reactions were performed as described in the culture dish; following quenching, cells were removed by trypsin and frozen as a dry pellet.

### Recovery of biotinylated proteins for mass spectrometry

Cell pellets were resuspended cold GdmCl lysis buffer containing 6 M Guanidinium Chloride (Sigma) and 100 mM Tris-HCl pH 8.5 (Sigma) containing 10 mM TCEP (Sigma) and 40 mM IAM (Sigma) and lysate was heated to 95 °C for 2x 5 mins followed by homogenization with a tip probe sonicator for 2x 20 sec cycles. Samples were diluted 1:1 in MilliQ H_2_O and precipitated O/N at - 20 °C with 4 volumes of acetone. Precipitated proteins were pelleted by centrifugation at 1,500 RPM for 5 min washed with 80% acetone at −20 °C, re-pelleted and air dried. Precipitates were resuspended in GdmCl Lysis buffer (without TCEP and IAM) and protein concentration was assayed with BCA kit (ThermoFisher) according to manufacturer’s protocol. 2 mg of total protein from each sample was diluted with MilliQ H_2_O to 1 M GdmCl in eqivolume. 1:2 (bead:protein (w:w)) washed streptavidin magnetic beads (Pierce) were added to each sample and incubated at 4 °C overnight at 1200 rpm on a thermomixer (ThermoFisher). The beads were washed 3×30 sec in GdmCl lysis buffer at 1200 rpm on the thermomixer at room temperature, followed by 2×30 sec in MilliQ H_2_O.

### TRF1 peptide synthesis and pull-down

N-terminus biotin labelled 21-25-mer TRF1 peptides were chemically synthesised by the CMRI peptide synthesis facility. Peptides were synthesised via solid phase peptide synthesis (SPPS) on a Syro II automated peptide synthesiser. Resin (50 mg, Chemmatrix rink amide, 0.36 mmol/g, 200-400 mesh) was washed (3 x 5 min in dimethylformamide (DMF)), swelled (DMF), deprotected with piperidine (20% in DMF, 2 x 1 min, 1 x 3 min, 1 x 10 min), washed again (3 x 5 min in DMF), and the first Fmoc protected amino acid coupled. Subsequent coupling steps were performed with standard Fmoc and side chain protected amino acids and chemistry, using 1,3-diisopropylcarbodiimide (3 eq) and Oxyma (3.3 eq) in DMF and heating where stable until reaching a phosphorylated residue or unstable amino acid. Fmoc-Ser(HPO3Bzl)-OH and Fmoc-Thr(HPO3Bzl)-OH were used to introduce the relevant phosphorylated amino acids and these couplings were performed at room temperature. Subsequent coupling steps were also performed at room temperature with double couples and extended coupling times. The final coupling of biotin to the N-terminus was performed manually under the same room temperature extended double coupling conditions. Cleavage and deprotection was performed using a standard cleavage cocktail of TFA:TIPS:Water (95:2.5:2.5). Peptides were precipitated in diethylether, redissolved in ACN:Water (30:70 with the addition of 0.1% FA to the final volume) and lyophilised. Peptides were then purified by reverse phase HPLC (Shimadzu Nexera, C16, A: Water, B: ACN, 5%-95% B over 45 minutes, 5 mL/min, 214 nm) and the target mass validated by MS (LCMS-2020, ESI-MS, [M+2H]2+ in the positive and [M-H]-in the negative). Purity was calculated by peak integration. Fractions were pooled and lyophilised affording the final product as a dry powder.

Lyophilised peptides were dissolved in 10 mM Tris-HCl pH 7.5 to a concentration of 1 mg mL^−1^ and bound to 0.6 mg of streptavidin magnetic beads (Pierce) for 2 hours at 4 °C. Mitotically arrested HT1080 6TG cells were lysed in immunoprecipitation buffer containing 20 mM HEPES pH 8 (Sigma), 150 mM KCl (Sigma), 0.5 mM EDTA (Sigma), 0.2% IGEPAL (Sigma), 0.5 mM DTT (Sigma) and 5% glycerol (Sigma), supplemented with PhosStop (Roche) and cOmplete Protease Inhibitor Cocktail (Roche). Cell lysate was mixed at 1:100 (v:v) with Benzonase (Merck) to digest precipitated chromatin. Lysate was precleared with fresh streptavidin magnetic beads and assayed by BCA (ThermoFisher) according to manufacturer’s instructions. 0.5 mg of total pre-cleared lysate was added to each peptide-streptavidin bead mix and shaken for 3 hours at 4 °C on a thermomixer at 1200 RPM. Non-specific interactors were removed by 2x washes with cold lysis buffer, followed by 3x washes with cold 100 mM ammonium bicarbonate solution to remove residual detergents. Samples were then processed as per proteomic sample preparation.

### Proteomic Sample Preparation

For APEX2/APEX-TRF1 samples, biotinylated proteins bound to streptavidin beads were resuspended in 100 mM ammonium bicarbonate (Sigma) and digested with 2 µg Trypsin (Promega) overnight at 37 °C shaking at 1200 RPM on a thermomixer. The supernatant of each sample was collected and acidified by adding up to 6 % trifluoroacetic acid (TFA) (Pierce) and 50% MS-grade acetonitrile (ACN) (Sigma). Precipitated lipids were removed by centrifugation at 14,000 RPM for 20 min, and the ACN was removed by vacuum centrifugation (GeneVac). Remaining supernatant was added to a house-packed stage-tip with 2x layers of C18 filter (Empore 3M) activated with ACN. Stage-tips were washed 2x with 0.1% TFA, eluted in 40% ACN /0.1% TFA into a thin-walled 96 well plate (ThermoFisher), and dried using vacuum centrifugation (GeneVac). Peptides were resuspended in MS buffer containing 2% ACN in 1% Formic Acid (Sigma).

For TRF1 peptide pulldown, proteins bound to streptavidin beads were digested in 50 mM ammonium bicarbonate and digested with 0.5 µg of trypsin overnight at 37 °C shaking at 1200 RPM on a thermomixer. Samples were resuspended in 2% TFA in isopropanol (IPA) (Sigma) and loaded on stage tips packed in-house with 2x layers of styrene-divinylbenzene (SDB-RPS) filter (AFFINISEP). Samples were washed with 1% TFA / 90% IPA and eluted in 5% ammonium hydroxide (Millipore) / 80% ACN into a thin walled 96 well plate. Desalted peptides were dried using vacuum centrifugation and resuspended in MS buffer.

### Liquid chromatography-Tandem mass spectrometry

Resuspended peptides were loaded into a Ultimate3000 UPLC with an autosampler maintained at 4 °C (Dionex), before loading onto a house-pulled 40 cm 75 µm I.D. fused silica column (Polymicro) containing 1.9 µm Reprosil AQ C18 particles (Dr. Maisch) in a 50 °C column oven (Sonation) on a nano-ESI source attached to a Q-Exactive Plus tandem mass spectrometer (ThermoFisher). APEX2/APEX2-TRF1 peptides were separated using the UPLC running a binary buffer system of A (0.1% Formic Acid) : B (90% ACN / 0.1% Formic Acid), over a gradient of 5-30% Buffer B over 150 min. An MS1 scan was acquired from 300-1600 m/Z (35,00 resolution, 3e6 AGC, 20 ms IT) followed by a data dependent MS2 scan with HCD dissociation in the Orbitrap (17,500 resolution, 1e5 AGC, 25 ms IT, 20 loop count). Thermo RAW files were acquired in centroid mode. TRF1 peptide pulldown samples were separated using the UPLC running a binary buffer system of A (0.1% Formic Acid) : B (90% ACN / 0.1% Formic Acid), over a gradient of 5-30% Buffer B over 50 min, followed by 30-60% Buffer B over 20 min. An MS1 scan was acquired from 300-1600 m/Z (35,00 resolution, 3e6 AGC, 20 ms IT) followed by a data dependent MS2 scan with HCD dissociation in the Orbitrap (17,500 resolution, 1e5 AGC, 25 ms IT, 20 loop count).

### Bioinformatic analysis

Thermo RAW files were processed with MaxQuant (v1.6.0.16) in standard settings, using a Human proteome database (Aug. 2018 release). Proteins were quantified using label-free quantification (LFQ) with additional identification enabled by match-between-run window set to 1.5 min. MaxQuant outputs were processed and analysed in Perseus (v1.6.10.43). Briefly, common contaminants, reverse database IDs and proteins IDs were removed. Datasets were filtered for proteins identified in < 5 (APEX2 samples) or < 3 (peptide pulldown samples) replicates of at least one sample group. Missing values of remaining proteins were imputed using the entire matrix (APEX2 samples were additionally batch corrected). To identify significantly enriched proteins, LFQ values for APEX2-TRF1 or phosphorylated peptides were tested against the respective APEX2 or non-phosphorylated peptide control samples using student’s t-test, filtering for p-value < 0.05. Gene ontology enrichment analysis was performed using Enrichr interactive webtool^72^. Initial comparisons of nocodazole and nocodazole + hesperadin datasets in the APEX2 samples revealed no significant differences in protein enrichment via streptavidin pulldown. To reduce the overall complexity of the analysis, only the nocodazole dataset has been analysed and presented.

### Metaphase-TIF, Interphase-TIF, and interphase streptavidin staining

For metaphase-TIF assays, mitotic chromosome spreads were obtained by cytospin as described previously^73^. Briefly, cells were collected by trypsinization and swelled in hypotonic buffer (0.2% KCl, 0.2% Tri-sodium citrate) for 10 min at room temperature. Cells were spread onto superfrost plus microscope slides (Epredia) at 2,000 rpm for 10 min by Cytospin 4 (Thermo Scientific) or Cellspin 1 (Tharmac). Samples were fixed in 2% paraformaldehyde (PFA)/1xPBS for 10 min. For interphase imaging, IMR90 E6E7 hTERT, HeLa, or MEFs were grown on 13 mm glass coverslips were fixed in 2% PFA/ 1x PBS for 10 min. Samples were incubated in KCM buffer (120 mM KCl, 20 mM NaCl, 10 mM Tris-HCl pH 7.5, 0.1% Triton X-100) for 10 min, followed by blocking in RNase A/ABDIL buffer (20 mM Tris-HCl pH 7.5, 2% BSA, 0.2% fish gelatin, 150 mM NaCl, 0.1% Triton X-100) at 37 °C for 30 min. For Interphase- and metaphase-TIF assays, cells were incubated with primary γ-H2AX antibody at 1:1,000 dilution at 37 °C for 1 hour. After 3 x 5 min washes in PBST buffer (1xPBS, 0.1% Tween-20) samples were incubates with secondary antibody. For interphase- and metaphase-TIF assays this was Alexa-568 goat anti-mouse IgG, 1:10,000 dilution at 37 °C for 30 min. For streptavidin staining we used streptavidin conjugated Alexa-488 at 1:1000 dilution at 37 °C for 60 min. Samples were washed in PBST for 5 min three times, then fixed in 2% PFA /1xPBS for 5 min. Samples were dehydrated sequentially in 70%, 95%, and 100% EtOH, then incubated with 0.3 ng mL^−1^ FAM-conjugated C-rich telomere peptide nucleic acid (PNA) probe (F1001, Panagene) in PNA buffer (70% formamide, 0.25% Blocking Reagent (NEN), 10mM Tris-HCl pH7.5, 5% MgCl_2_ buffer (82 mM Na_2_HPO_4_, 9 mM citric acid, 25 mM MgCl_2_)) at 80 °C for 8 min. Following overnight incubation at 37 °C, samples were washed in PNA Wash A buffer (70% formamide, 10 mM Tris-HCl pH7.5) for 15 min twice and PNA Wash B buffer (50 mM Tris-HCl pH7.5, 150 mM NaCl, 0.8% Tween-20) for 10 min three times (0.2 µg mL^−1^ DAPI in the second wash) and mounted with Vectashield PLUS Antifade (H-2000, Vector Laboratories) or ProLong Gold (Life Technologies).

For IMR90 E6E7 hTERT fibroblasts, images of interphase nuclei and metaphase spreads were taken with a 100x objective lens (PlanApo/1.45-NA oil) on a BZ-X710 microscope (KEYENCE) and analyzed by automated counting with Hybrid Cell Count and Macro Cell Count within the BZ-X Analyzer software (KEYENCE). For MEF metaphase-TIF experiments, images were captured using a ZEISS AxioImager Z.2 with a 63x, 1.4NA oil objective and appropriate filter cubes, using a CoolCube1 camera (Metaystems). Automated metaphase finding and image acquisition for these experiments were done using the MetaSystems imaging platform as described elsewhere^29^. For imaging of HeLa and MEF interphase cells, images were captured using Zen software and a ZEISS AxioImagerZ.2, with a 63x, 1.4NA oil objective, appropriate filter cubes and an Axiocam506 monochromatic camera (ZEISS).

### Telomere fluorescence intensity measurements

To compare telomere FISH signal between sister chromatids, identifiable sister chromatid pairs were chosen from metaphase-TIF images and telomere signals were manually quantified using the oval selection tool within ImageJ2 software (2.14.0/1.54h). For each sister telomere pair, an ROI of minimal size was established, and the signals at the telomeres and in the background region between sister telomeres were measured as an average signal within the ROI. The background signal was deducted from the telomere signals before calculating signal ratio within each sister telomere pair and plotting as the log_2_ value.

### Telomere-loop fluorescence imaging

Samples were prepared for super-resolution imaging of telomere macromolecular structure as described elsewhere^11^ with slight modification. Briefly, mitotic cells were pelleted at 1,000 g for 5 min at 4 °C and washed with ice-cold nuclei wash buffer (10 mM Tris-HCl [pH 7.4], 15 mM NaCl, 60 mM KCl, 5 mM EDTA, 300 mM sucrose). The sample was re-suspended in nuclei wash buffer in a 6-well non-tissue culture treated plate and incubated for 5 min while stirring on ice, in the dark, in the presence of 100 mg mL^−1^ Trioxsalen (Sigma) diluted in DMSO. The material was then exposed to 365 nm UV light, 2-3 cm from the light source (model UVL-56, UVP), for 30 min, while incubated on ice with continuous stirring. After cross-linking, the material was centrifuged at 1000 g for 5 min, washed with ice-cold nuclei wash buffer. Pre-warmed 37 °C spreading buffer (10 mM Tris-HCl [pH 7.4], 10 mM EDTA, 0.05% SDS, 1 M NaCl) was added to the sample and deposited on a 18 x 18 mm 170 µm thick coverslip using a Cellspin1 (Tharmac) at 1000 rpm for 1 min. Coverslips were fixed in −20 °C methanol for 10 min followed by 1 min in −20 °C acetone. Coverslips were rinsed with PBS then dehydrated through a 70%, 95%, and 100% ethanol series. Ethanol dehydrated coverslips were denatured for 10 min at 80 °C in the presence of C-rich telomere PNA probe conjugated to Alexa fluor 488 (Alexa488-OO-ccctaaccctaaccctaa, Panagene, F1004). PNA probe was prepared and diluted to 0.3 ng mL^−1^ as described previously^41^. Following hybridization overnight in a dark humidified box, coverslips were washed twice in PNA Wash A (70% Formamide; 10 mM Tris-HCL [pH 7.5]) and thrice in PNA Wash B (50 mM Tris-HCL [pH 7.5], 150 mM NaCl, 0.8% Tween-20) with gentle shaking. DAPI was added in the second wash of PNA wash B to stain the chromatin. Coverslips were rinsed in MilliQ water and dehydrated through a 70%, 95%, and 100% ethanol series. Slides were mounted in Prolong Gold (Life Technologies) in the presence of DAPI. Airyscan imaging was performed on a ZEISS LSM880 AxioObserver confocal fluorescent microscope fitted with an Airyscan detector and a Plan-Apochromat 633 1.4 NA M27 oil objective. Alexa Fluor 488 labeled telomeres were captured with 4% excitation power of 488 nm laser, 131 binning, detector gain of 950 and digital gain of 1 in super-resolution mode. A total of 5 z stacks (200 nm) were captured with frame scanning mode, unidirectional scanning, and line averaging of 2 in 1024 x 1024 pixels at 89.88 mm x 89.88 mm to scale. Z stacks were Airyscan processed using batch mode in Zen Black software (ZEISS). Images were maximum intensity projected and scored by eye with the researcher blinded to experimental conditions.

### Live cell imaging and analysis

Cell Observer inverted wide field microscope (Zeiss) with 20 × 0.8 NA air objective was utilized to perform differential interference contrast imaging to visualise mitotic duration and outcomes. Cells were cultured on a glass bottom 24-well plate (MatTek Corporation) at 37 °C, 10% CO_2_ and atmospheric oxygen provided by Zeiss Incubation System Module CELLS. Cells were on the microscope for 3 hr to achieve stability in the focal plane and images were captured every 6 min up to 63 hr using an Axiocam 506 monochromatic camera (Zeiss) and Zen software (Zeiss). For all movies, mitotic duration was scored by eye and calculated from nuclear envelope breakdown until cytokinesis or mitotic death. The imaging analysis was performed using Zen software.

### Immunoprecipitation

Trypsinised HT1080-6TG cells were resuspended in cold non-denaturing lysis buffer containing 20 mM Tris-HCl pH 8.0, 137 mM NaCl, 1% IGEPAL, 2 mM EDTA supplemented with PhosStop and cOmplete Protease Inhibitor cocktail. Nuclei were mechanically dissociated through a 27 G syringe before removing insoluble chromatin by centrifugation at 14,000 RPM. Protein concentration was determined by BCA per the manufacturer’s instructions and precleared with 50 µg of protein A/G magnetic beads (Peirce) for 30 min at 4 C mixing at 1200 RPM. Washed protein A/G beads were incubated with primary antibody in non-denaturing lysis buffer containing 100 µg mL^−1^ BSA for 1 hour at 4 °C on a mixer at 1200 RPM. 0.1 - 0.2 mg of antibody bound beads were added to 0.5 - 2 mg of total protein lysate and shaken for 2 hours at 4 °C mixing at 1200 rpm (for anti-TRF1 pulldowns, bead incubation was increased to overnight, on a nutator). Non-specific interactors were removed by 3×1 min washes with IP wash buffer containing 20 mM Tris-HCl pH 8.0, 300 mM NaCl, 1% IGEPAL, and 2 mM EDTA. Proteins were eluted by mixing in 2xLDS buffer (-EDTA) containing β-mercaptoethanol at RT for 5 mins. Immunoprecipitations were analysed by western blotting, comparing to 0.5-1% (w:w) of input lysate.

### In vitro kinase assay

Thymidine synchronised HT1080 6TG cells were processed as per immunoprecipitation protocol using 80 µg of total lysate per kinase reaction, incubating overnight with antibody-bead mix. Following removal of non-specific interactors with IP wash buffer, beads were rinsed with kinase assay buffer containing 25 mM MOPS pH 7.2, 25 mM MgCl_2_, 5 mM EGTA, 5 mM EDTA and 0.25 mM DTT diluted 1:5 in a sterile 50 µg mL^−1^ BSA solution. Beads were resuspended in diluted kinase assay buffer containing 0.5 mM ATP and recombinant Aurora B kinase (AbCam). Kinase reactions were shaken at 30 °C for 10 mins at 1000 RPM and quenched with 4x LDS buffer (-EDTA) containing β-mercaptoethanol. Assays were analysed and detected per western blotting protocol. Resultant immunoblots were analysed by densitometry corrected for background and loading against total TRF1 or 2. Enzyme kinetics were estimated using non-linear Michaelis-Menten fitting in Prism, reporting K_m_ within a 95% confidence interval.

### Western blotting

For IMR90 E6E7 hTERT fibroblasts, whole-cell extracts were resolved using 4-20% Mini-PROTEAN TGX precast gels (BioRad) and transferred to PVDF membranes (Millipore). Membranes were blocked with Blocking Buffer (Nacalai) and incubated with primary antibodies followed by incubation with secondary antibodies. Signal developed by Chemi-Lumi Super One (Nacalai) was detected by LAS-3000 imager (Fujifilm). For HeLa, HT1080 6TG, and MEFs whole cell extracts were prepared by lysing cells in 4xLDS buffer (-EDTA) containing β-mercaptoethanol and 500 U Benzonase (Millipore) at 10,000 cells / µL for 10-15 min at RT, followed by denaturation at 65 °C for 10 min. Proteins were resolved using precast 1.5 mm 4-12% Bis-Tris NuPage gels (ThermoFisher) and transferred to 0.2 µm nitrocellulose membranes (GEHE10600008, Amersham). For total proteins, membranes were blocked in 5% skim milk powder / 1xTBS containing 0.1% Tween-20 (TBS-T), for phospho-specific antibodies membranes were blocked in 5% BSA / TBS-T for 1 hour at RT with gentle shaking, for total biotin blots membranes were blocked overnight in 3% BSA / TBS-T without primary antibodies. Membranes were incubated in primary antibodies diluted in 5% BSA / TBS-T overnight followed by incubating with secondary antibodies in 5% skim milk powder / TBS-T for 1 hour at room temperature. Following block, total biotin blots were treated with streptavidin-HRP at a concentration 0.3 µg mL^−1^ in 3% BSA /TBS-T for 1 hour at RT. HRP signal was detected in 9:1 mix of Clarity:Clarity Max ECl reagent (BioRad) and captured using a Chemidoc Touch imager (BioRad).

### Antibodies and recombinant proteins

Antibodies and their dilution for Western blotting were as follows: TRF2 (NB110-57130SS, Novus Biologicals), 1:1000; TRF1 (sc-56807, SantaCruz), 1:1000; TRF1 rabbit polyclonal antibody (Ishikawa lab, used in Extended data Figure 3), 1:1,000; BLM (NB100-214, NovusBio), 1:5000; Flag (F1804, Sigma), 1:2000; Myc (9B11 Cell Signalling Technology), 1:1000; Top3A (14525-1-AP, Proteintech), 0.3 µg mL^−1^; StreptAvidin-HRP (S911, ThermoFisher), 1:1000; INCENP (ab-12183, AbCam), 1:2000; Aurora B (ab-2254, AbCam), 1:1000; Borealin (ab74473, Abcam), 1:1000; Survivin (NB500-201, Novus Biologicals), 1:1000; beta-Actin (A5441, Sigma), 1:20,000; Actin (MAB1501R, Millipore), 1:10,000; Vinculin (V9131, Sigma), 1:5000; GAPDH (MAB374, Millipore), 1:5000; Goat anti-mouse HRP (P044701, DAKO Agilent), 1:5000; and Goat anti-Rabbit HRP (P044801, DAKO Agilent). For immunofluorescence we used: γ-H2AX (Clone 2F3, Biolegend) 1:200 or (05-636, Merck Millipore), 1:500; Alexa-568 goat anti-mouse (A11031, Invitrogen); 1:10,000; and Streptavidin Alexa-488 (S32354, Invitrogen), 1:1,000.

Polyclonal rabbit antibodies against phopsho-TRF1 and TRF2 were generated by Genscript. Briefly, animals were immunized with peptides corresponding to pTRF1-T358 (SRRA(p)TESRIPVSKS), and pTRF2-Ser65 (ASRS(p)SGRARRGRHEC). Antibodies were purified by antigen affinity purification selecting for minimized cross adsorption and validated using indirect ELISA. For western blotting, antibodies were diluted in 5% BSA / TBS-T at a concentration of 0.2 µg mL^−1^.

### Statistical analysis and figure preparation

GraphPad Prism 9 was used for all the statistical analysis for this study. The number of experimental replicates is indicated in the figure legends. Comparisons between two groups were performed using Mann–Whitney test. Multiple comparisons were performed using Kruskal-Wallis followed by Dunn’s test unless otherwise indicated in the figure legends. For metaphase-TIF analysis in IMR90 E6E7 hTERT fibroblasts, images were taken blindly by lab technicians and quantified using automated BZ-X Analyzer software. For metaphase-TIF analysis in all other cell lines, and for the analysis of telomere macromolecular structure, scientists were blinded to the experimental conditions. Figures were prepared using Adobe Illustrator and Affinity Publisher.

### Data Availability

The mass spectrometry outputs, and MaxQuant analyses in this paper will be made available on the PRIDE database following final publication. The proteomics and other datasets generated and/or analysed during the current study are available from M.T.H. and A.J.C. upon reasonable request.

## SUPPLEMENTARY FIGURES AND LEGENDS

**Figure S1.**
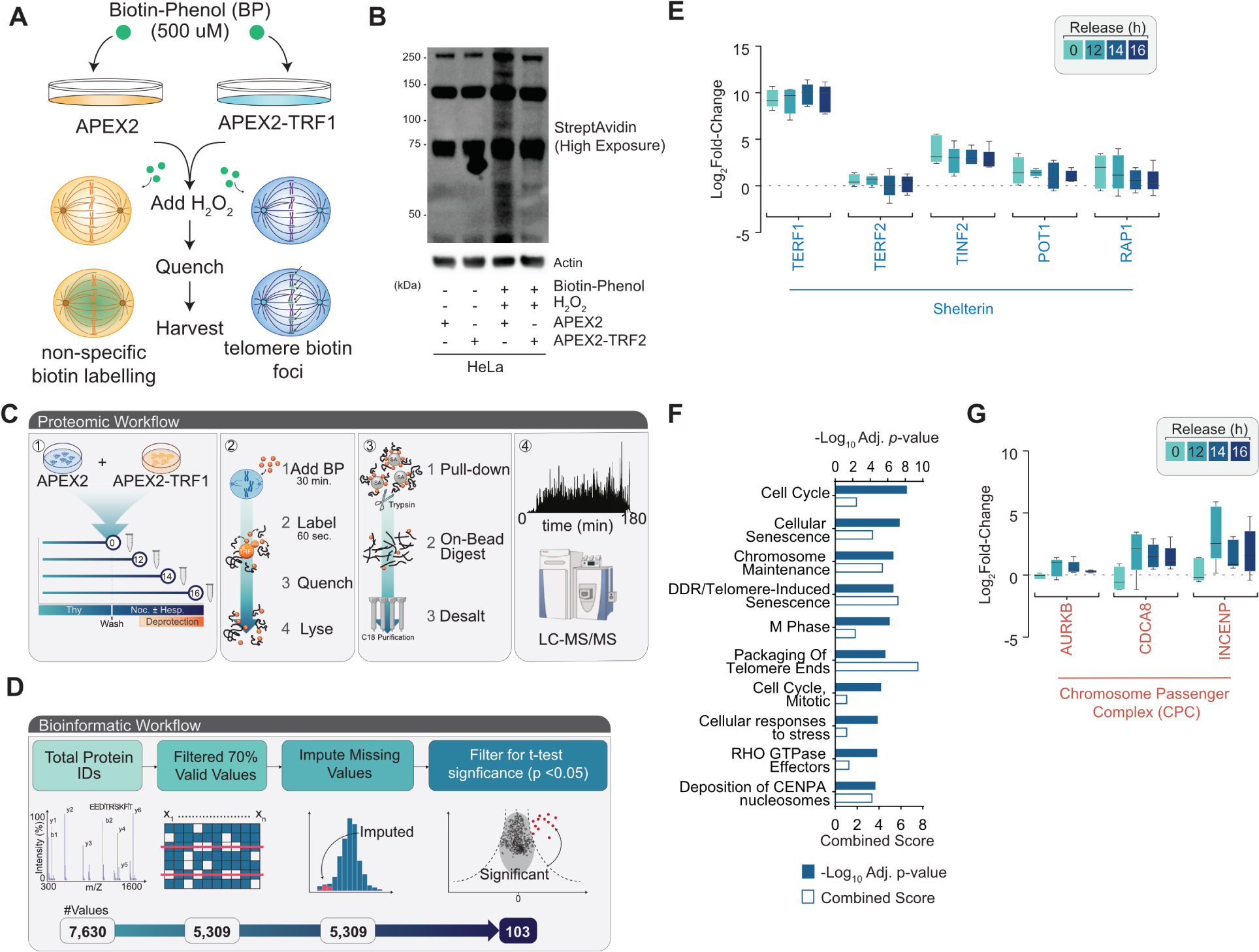
Proximity dependent TRF1 interactomics reveals components of the MAD telomere deprotection pathway, related to Figure 1. **A.** Schematic of APEX2-TRF1 proximal biotin-labelling for interactomics. **B.** Immunoblot of streptavidin bound to whole cell extracts derived from APEX2 or APEX2-TRF1 HeLa cells with or without APEX2 activation with Biotin-phenol and hydrogen peroxide (H_2_O_2_) (representative example of n = 3 biological replicates). **C, D.** Schematic of the proteomics (C) and bioinformatics (D) workflow for APEX2-TRF1 interactomics. **E.** Log_2_-fold change in streptavidin-precipitated shelterin proteins between APEX2-TRF1 and APEX2 samples (Tukey’s boxplot, median ± IQR, n = 5 biological replicates). **F.** Enrichr gene ontology analysis of proteins significantly enriched within mitotic Flag-APEX2-TRF1 datasets. Ontology terms are ranked by –Log_10_ p-value (filled bars), reporting Enrichr combined score (empty bars). G. Log_2_-fold change in streptavadin-precipitated Chromosome Passenger Complex proteins between APEX2-TRF1 and APEX2 samples (Tukey’s boxplot, median ± IQR, n = 5 biological replicates).

**Figure S2.**
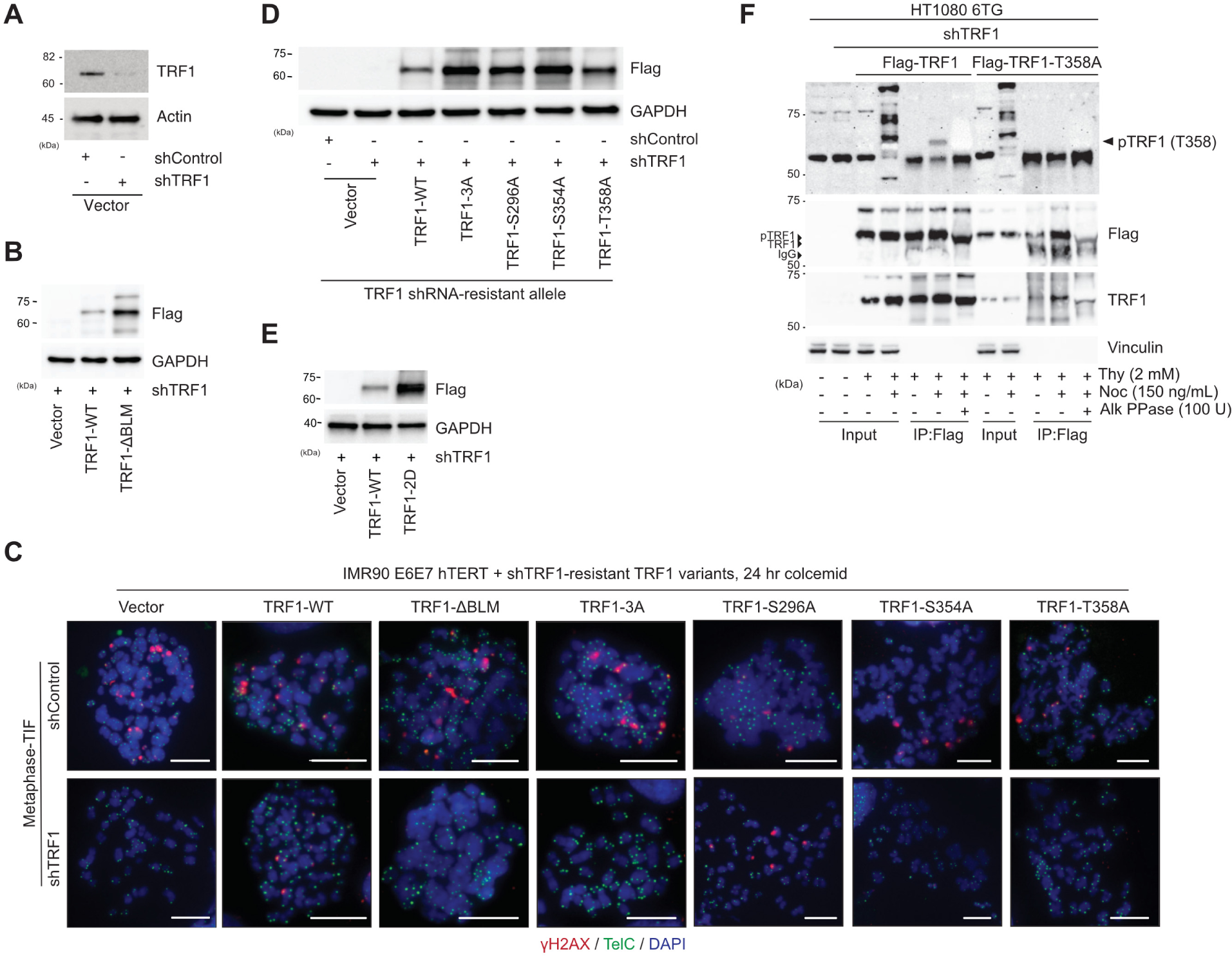
AURKB consensus sites at TRF1-S354 and -T358 participate in MAD-telomere deprotection. **A.** Immunoblots of whole cell extracts derived from Control or TRF1 shRNA IMR90 E6E7 hTERT fibroblasts (representative example of n ≥ 3 biological replicates). **B.** Immunoblots of whole cell extracts derived from Control or TRF1 shRNA IMR90 E6E7 hTERT fibroblasts expressing the indicated shRNA resistant Flag-TRF1 alleles (representative example of n ≥ 3 biological replicates). **C.** Metaphase-TIF assays using combined γ-H2AX immunofluorescence (red) and telomere FISH (green) in control or TRF1 shRNA IMR90 E6E7 hTERT cultures expressing the indicated shRNA-resistant FLAG-TRF1 alleles. All images are from cells treated with 100 ng mL^−1^ colcemid for 24 hours prior to sample collection (representative images from n ≥ 3 biological replicates). Scale bar, 10µm, the DNA is stained with DAPI (blue). **E.** Immunoblots of whole cell extracts derived from control or TRF1 shRNA IMR90 E6E7 hTERT fibroblasts expressing the indicated shRNA resistant TRF1 alleles (representative example of n = 3 biological replicates). **F.** Immunoblots of Flag immuno-precipitates from TRF1 shRNA HT1080 6TG cells expressing shRNA-resistant WT Flag-TRF1 or Flag-TRF1-T358A. Where indicated, cells were synchronised with a double-thymidine block (Thy) and released in the presence or absence of 150 ng mL^−1^ nocodazole (Noc) for 16 hours before sample collection. Where indicated extracts were treated with alkaline phosphatase (Alk. PPase). The pTRF1-T358 band is indicated with an arrow. Representative example of n = 3 biological replicates is shown.

**Figure S3.**
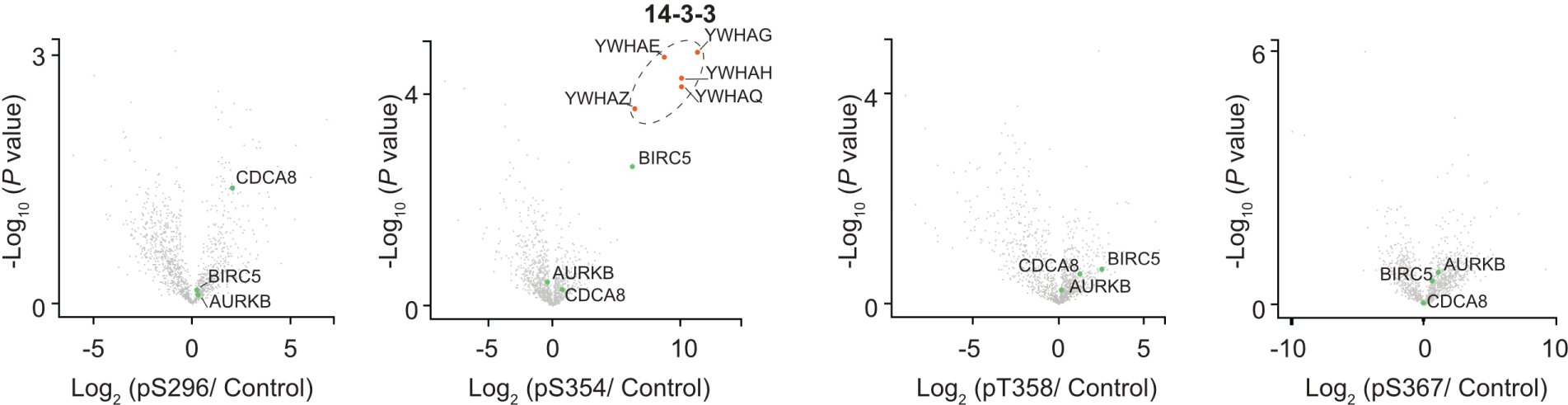
TRF1 phospho-peptides pull down Survivin, related to Figure 3. Volcano plots representative of LC-MS/MS analysis of TRF1 peptide pulldowns as depicted in Fig. 3A. LFQ values from proteins detected in phosphorylated TRF1 peptide pulldowns were compared to respective values in non-phosphorylated controls and plotted as a function of Log_2_ fold-change and –Log_10_ p-value, indicating enrichment of detected CPC members, and 14-3-3 phospho-binding protein (student’s T-test of n=3 biological replicates).

**Figure S4.**
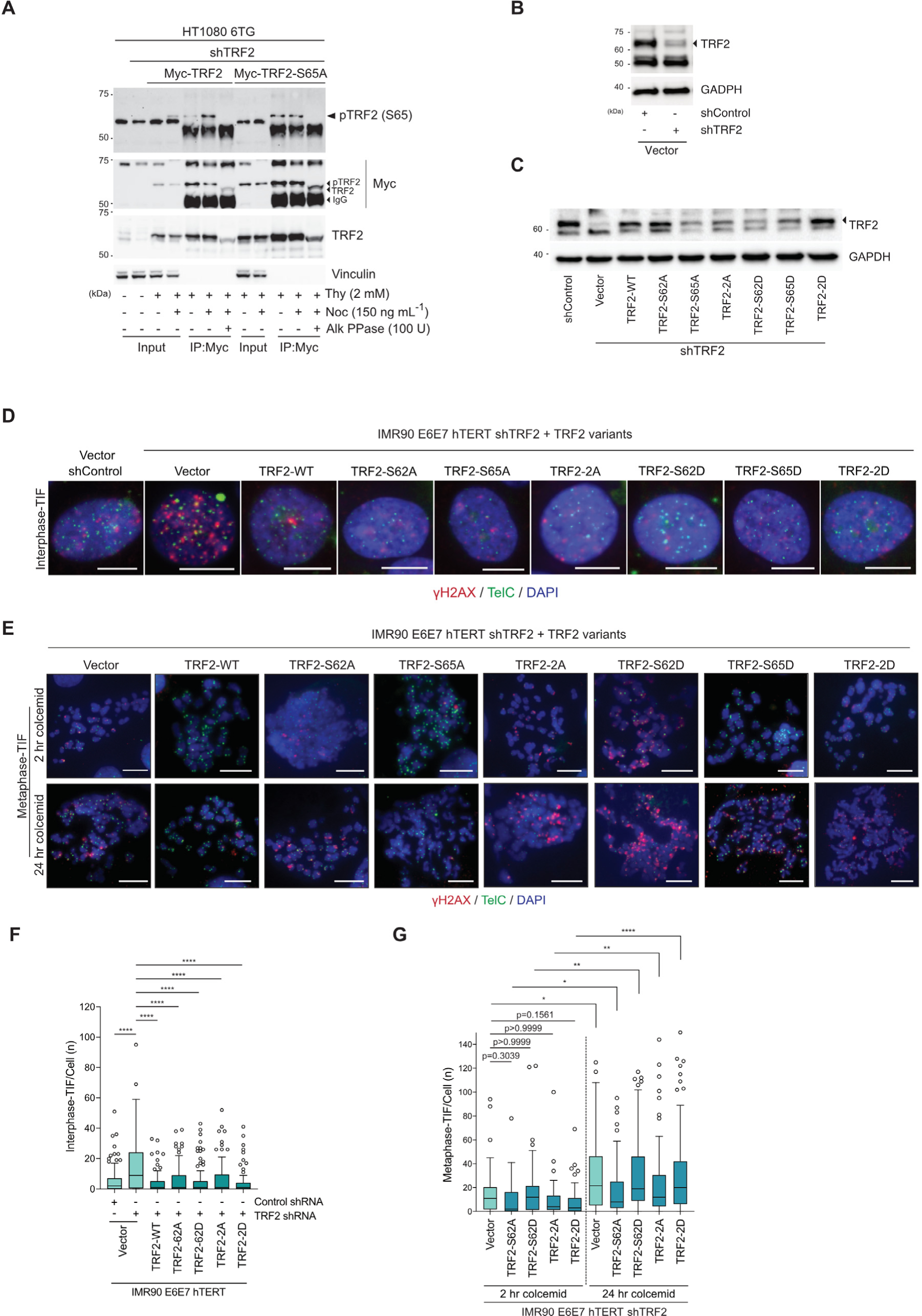
The TRF2 basic domain functions in mitosis-specific telomere protection, related to Figure 4. **A.** Immunoblots of Myc immuno-precipitates from TRF2 shRNA HT1080 6TG cells expressing WT Myc-TRF2 or Myc-TRF2-S65A. Where indicated, cells were synchronised with a double-thymidine block (Thy) and released in the presence or absence of 150 ng mL^−1^ Nocodazole (Noc) for 16 hours before sample collection. Where indicated extracts were treated with alkaline phosphatase (Alk. PPase). Representative example of n = 3 biological replicates is shown. **B.** Western blots of whole cell extracts derived from IMR90 E6E7 hTERT TRF2 tranduced with Control or TRF2 shRNA (representative example of n ≥ 3 biological replicates). **C.** Western blots of whole cell extracts derived from IMR90 E6E7 hTERT transduced with TRF2 shRNA and the indicated Myc-TRF2 alleles (representative example of n ≥ 3 biological replicates). **D, E.** Examples of Interphase-TIF (D) and Metaphase-TIF (E) assays in IMR90 E6E7 hTERT cells transduced with the indicated shRNA and TRF2 alleles (representative example of n ≥ 3 biological replicates). In (E) cultures were treated with 100 ng mL^−1^ colcemid for 2 or 24 hours prior to sample collection **F.** Interphase-TIF in TRF2 shRNA IMR90 E6E7 hTERT fibroblasts expressing the indicated TRF2 alleles (n = 3 biological replicates of 45 nuclei per replicate, pooled into a Tukey box plot, Kruskal-Wallis followed by Dunn’s multiple comparisons test, ****p<0.0001). **G.** Metaphase-TIF in TRF2 shRNA IMR90 E6E7 hTERT fibroblasts expressing the indicated TRF2 alleles. Cells were treated with 2 or 24 hours of 100 ng mL^−1^ colcemid prior to sample collection (n = 3 biological replicates of 15 and 30 metaphases per replicate for 2 hours and 24 hours colcemid, respectively, pooled into a Tukey box plot, Kruskal-Wallis followed by Dunn’s multiple comparisons test, ****p<0.0001).

**Figure S5:**
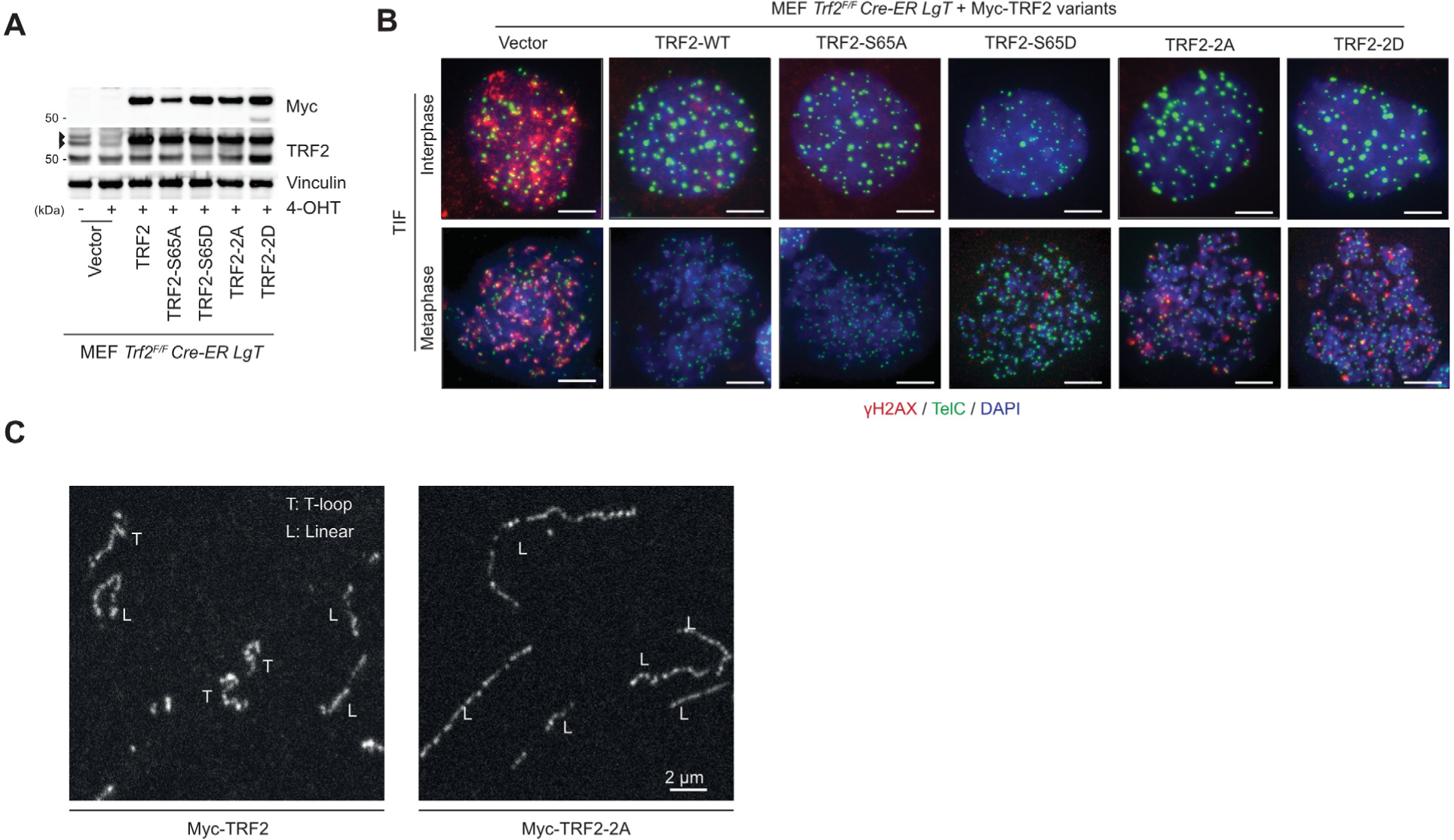
Imaging telomere deprotection and telomere macromolecular structure in *Trf2^F/F^ Cre-ER LgT* MEFs, related to Figure 5. **A.** Representative examples of immunoblots of whole cell extracts derived from *Trf2^F/F^ Cre-ER LgT* MEFs expressing the indicated Myc-tagged hTRF2 alleles. Representative examples of immunoblots of whole cell extracts derived from *Trf2^F/F-^ Cre-ER LgT* MEFs expressing the indicated Myc-tagged hTRF2 alleles. Where indicated the cultures were treated with 4-OHT for 36 hours prior to sample collection to delete endogenous *mTrf2*. **B.** Representative interphase-TIF and metaphase-TIF images from *Trf2^F/F^ Cre-ER LgT* MEFs expressing hTRF2-WT or the indicated hTRF2 variants. Representative interphase-TIF and metaphase-TIF images from *Trf2^F/F-^ Cre-ER LgT* MEFs expressing hTRF2-WT or the indicated hTRF2 variants. In all conditions endogenous *mTrf2* was deleted by 4-OHT addition 36 hours prior to sample fixation. Samples were stained with combined γ-H2AX immunofluorescence (red) and telomere FISH (green). The DNA is labelled with DAPI (blue). For Metaphase-TIF cells were treated with 400 ng mL^−1^ Nocodazle for 14 hours prior to sample collection. Scale bar, 10µm. Representative example of n = 3 biological replicates is shown. **C.** Representative fields from AiryScan microscopy of telomere macromolecular structure. *Trf2^F/F^ Cre-ER LgT* MEFs expressing Myc-TRF2-WT or Myc-TRF2-2A (S62A and S65A) were treated with 4-OHT for 36 hours prior to sample collection to delete endogenous *Trf2*. *Trf2^F/F^ Cre-ER LgT* MEFs expressing Myc-TRF2-WT or Myc-TRF2-2A (S62A and S65A) were treated with 4-OHT for 36 hours prior to sample collection to delete endogenous *Trf2*. Cultures were treated with 400 ng mL^−1^ Nocodazle for 14 hours and collected by mitotic shake-off before trioxsalen cross-linked in situ, chromatin spreading on coverslips through cytocentrifugation, and telomere FISH staining. Linear (L) and t-loops (T) are shown. Scale bar, 2 µm. Representative example of n = 3 biological replicates is shown.

**Figure S6.**
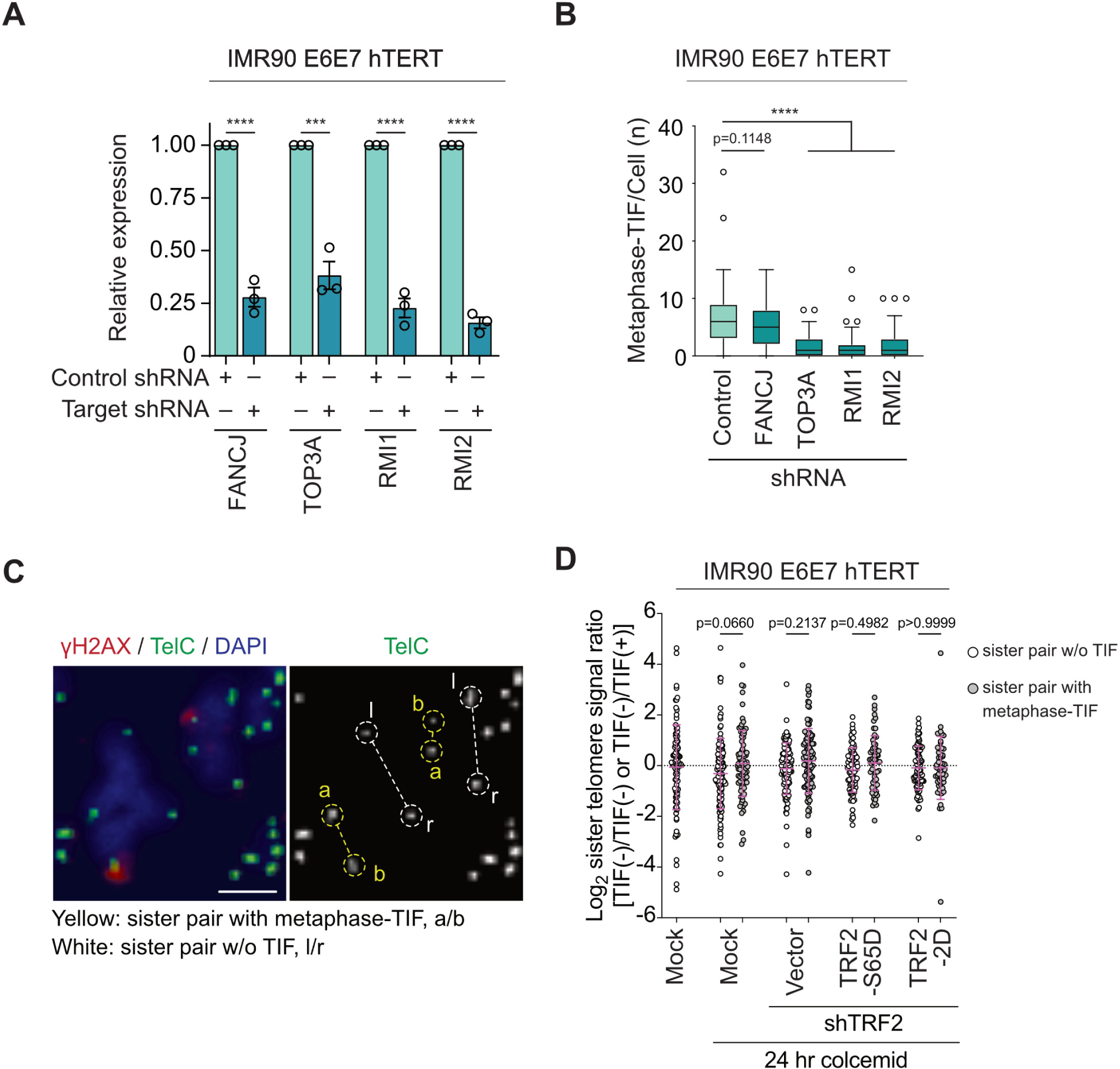
BTR promotes MAD telomere deprotection without telomere shortening, related to Figure 6. **A.** Relative gene expression in shRNA transduced IMR90 E6E7 hTERT. For each targeted gene indicated below, transcription was measured by RT-qPCR and normalized to Actin (mean +/− s.e.m., n = 3 biological replicates, Unpaired two-tailed t test, ****p<0.0001). **B.** Metaphase-TIF in IMR90 E6E7 hTERT fibroblasts five days post-transduction with the indicated shRNAs. Cultures were treated with 100 ng mL^−1^ colcemid for 24 hours prior to sample collection (n = 3 biological replicates of 30 metaphases per replicate, compiled into a Tukey Box plot, Kruskal-Wallis followed by Dunn’s multiple comparisons test, ****p<0.0001). **C.** Measurement of inter-chromatid telomere length variability. On metaphase-TIF assays stained with γ-H2AX IF (red), telomere FISH (TelC, Green), and DAPI, we identified sister telomeres without γ-H2AX staining (sister pair without (w/o) metaphase-TIF), or where a single chromatid was γ-H2AX positive (sister pair with metaphase-TIF). Telomere length for each chromatid was measured by FISH intensity. We calculated the ratio between telomere lengths of the sister chromatids and plotted as log_2_ values. **D.** Ratio of sister telomere lengths as described in (C) for mock or TRF2 shRNA transduced IMR90 E6E7 hTERT cells ± the indicated exogenous TRF2 alleles. Where indicated cultures were treated with 100 ng mL^−1^ colcemid for 24 hours prior to sample collection (mean ± s.d., n ≥ 54, ordinary one-way ANOVA followed by Šídák’s multiple comparisons test, F=2.048, DF=(7, 696)).

**Figure S7.**
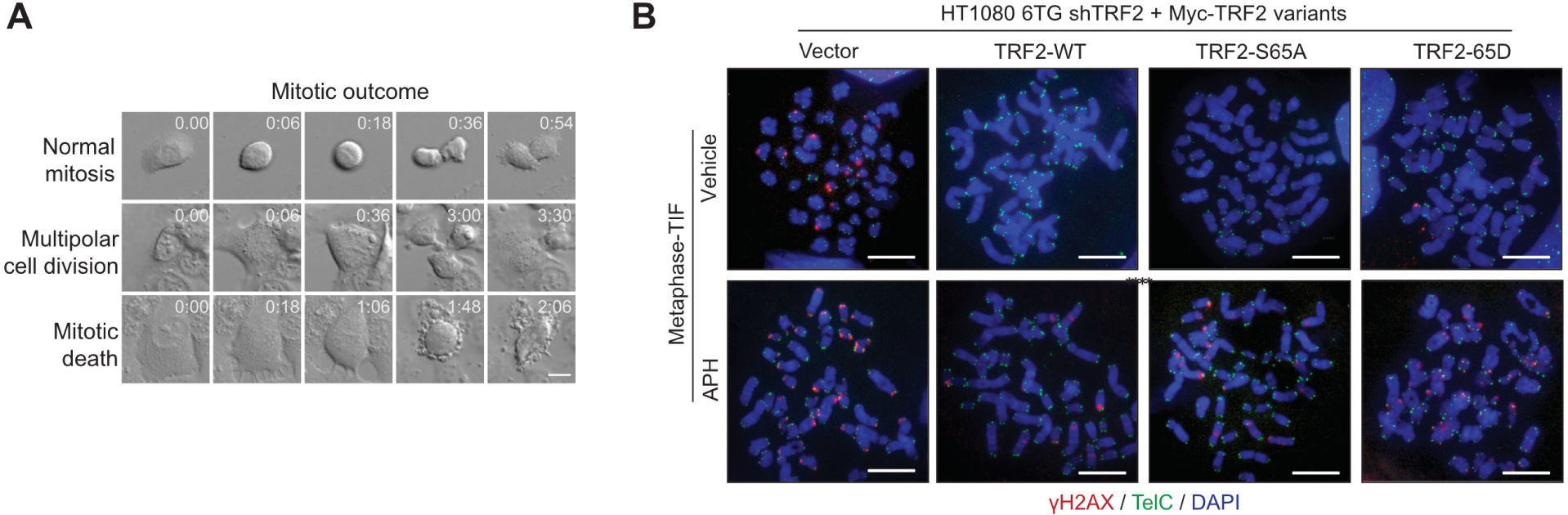
Imaging mitotic death and telomere deprotection in cells experiencing physiological mitotic arrest, related to Figure 7. **A.** Captures from differential interference contrast live imaging of HT1080 6TG cells treated with 1µM Aphidicolin (APH). Mitotic outcomes are indicated. Time is shown as hours: minutes relative to the first images. Scale bar, 20 µm. Representative example of n = 3 biological replicates. **B.** Representative metaphase-TIF images from TRF2 shRNA HT1080 6TG cultures expressing the indicated Myc-TRF2 alleles. Cultures were treated with 1µM Aphidicolin (APH) for 40 hours before sample collection Scale bar, 10µm. Representative example of n = 3 biological replicates.

## Notes

### Competing Interest Statement

The authors have declared no competing interest.

## REFERENCES

1. de Lange, T. (2018). Shelterin-Mediated Telomere Protection. Annual review of genetics 52, 223–247. 10.1146/annurev-genet-032918-021921.

2. Cesare, A.J., and Karlseder, J. (2012). A three-state model of telomere control over human proliferative boundaries. Current opinion in cell biology 24, 731–738. 10.1016/j.ceb.2012.08.007.

3. Karlseder, J., Smogorzewska, A., and de Lange, T. (2002). Senescence induced by altered telomere state, not telomere loss. Science (New York, N.Y 295, 2446–2449.

4. d’Adda di Fagagna, F., Reaper, P.M., Clay-Farrace, L., Fiegler, H., Carr, P., Von Zglinicki, T., Saretzki, G., Carter, N.P., and Jackson, S.P. (2003). A DNA damage checkpoint response in telomere-initiated senescence. Nature 426, 194–198.

5. Kaul, Z., Cesare, A.J., Huschtscha, L.I., Neumann, A.A., and Reddel, R.R. (2011). Five dysfunctional telomeres predict onset of senescence in human cells. EMBO reports 13, 52–59. 10.1038/embor.2011.227.

6. Griffith, J.D., Comeau, L., Rosenfield, S., Stansel, R.M., Bianchi, A., Moss, H., and de Lange, T. (1999). Mammalian telomeres end in a large duplex loop. Cell 97, 503–514.

7. Munoz-Jordan, J.L., Cross, G.A., de Lange, T., and Griffith, J.D. (2001). t-loops at trypanosome telomeres. The EMBO journal 20, 579–588.

8. Cesare, A.J., Quinney, N., Willcox, S., Subramanian, D., and Griffith, J.D. (2003). Telomere looping in P. sativum (common garden pea). Plant J 36, 271–279. 10.1046/j.1365-313x.2003.01882.x.

9. Yu, Q., Gates, P.B., Rogers, S., Mikicic, I., Elewa, A., Salomon, F., Lachnit, M., Caldarelli, A., Flores-Rodriguez, N., Cesare, A.J., et al. (2022). Telomerase-independent maintenance of telomere length in a vertebrate. bioRxiv, 2022.2003.2025.485759. 10.1101/2022.03.25.485759.

10. Nikitina, T., and Woodcock, C.L. (2004). Closed chromatin loops at the ends of chromosomes. The Journal of cell biology 166, 161–165.

11. Van Ly, D., Low, R.R.J., Frolich, S., Bartolec, T.K., Kafer, G.R., Pickett, H.A., Gaus, K., and Cesare, A.J. (2018). Telomere Loop Dynamics in Chromosome End Protection. Molecular cell 71, 510–525 e516. 10.1016/j.molcel.2018.06.025.

12. Doksani, Y., Wu, J.Y., de Lange, T., and Zhuang, X. (2013). Super-resolution fluorescence imaging of telomeres reveals TRF2-dependent T-loop formation. Cell 155, 345–356. 10.1016/j.cell.2013.09.048.

13. Okamoto, K., Bartocci, C., Ouzounov, I., Diedrich, J.K., Yates, J.R., 3rd, and Denchi, E.L. (2013). A two-step mechanism for TRF2-mediated chromosome-end protection. Nature 494, 502–505. 10.1038/nature11873.

14. Benarroch-Popivker, D., Pisano, S., Mendez-Bermudez, A., Lototska, L., Kaur, P., Bauwens, S., Djerbi, N., Latrick, C.M., Fraisier, V., Pei, B., et al. (2016). TRF2-Mediated Control of Telomere DNA Topology as a Mechanism for Chromosome-End Protection. Molecular cell 61, 274–286. 10.1016/j.molcel.2015.12.009.

15. Ruis, P., Van Ly, D., Borel, V., Kafer, G.R., McCarthy, A., Howell, S., Blassberg, R., Snijders, A.P., Briscoe, J., Niakan, K.K., et al. (2021). TRF2-independent chromosome end protection during pluripotency. Nature 589, 103–109. 10.1038/s41586-020-2960-y.

16. Markiewicz-Potoczny, M., Lobanova, A., Loeb, A.M., Kirak, O., Olbrich, T., Ruiz, S., and Lazzerini Denchi, E. (2021). TRF2-mediated telomere protection is dispensable in pluripotent stem cells. Nature 589, 110–115. 10.1038/s41586-020-2959-4.

17. Herbig, U., Jobling, W.A., Chen, B.P., Chen, D.J., and Sedivy, J.M. (2004). Telomere shortening triggers senescence of human cells through a pathway involving ATM, p53, and p21(CIP1), but not p16(INK4a). Molecular cell 14, 501–513.

18. Chin, L., Artandi, S.E., Shen, Q., Tam, A., Lee, S.L., Gottlieb, G.J., Greider, C.W., and DePinho, R.A. (1999). p53 deficiency rescues the adverse effects of telomere loss and cooperates with telomere dysfunction to accelerate carcinogenesis. Cell 97, 527–538.

19. Counter, C.M., Avilion, A.A., LeFeuvre, C.E., Stewart, N.G., Greider, C.W., Harley, C.B., and Bacchetti, S. (1992). Telomere shortening associated with chromosome instability is arrested in immortal cells which express telomerase activity. The EMBO journal 11, 1921–1929.

20. Nassour, J., Radford, R., Correia, A., Fuste, J.M., Schoell, B., Jauch, A., Shaw, R.J., and Karlseder, J. (2019). Autophagic cell death restricts chromosomal instability during replicative crisis. Nature 565, 659–663. 10.1038/s41586-019-0885-0.

21. Nassour, J., Aguiar, L.G., Correia, A., Schmidt, T.T., Mainz, L., Przetocka, S., Haggblom, C., Tadepalle, N., Williams, A., Shokhirev, M.N., et al. (2023). Telomere-to-mitochondria signalling by ZBP1 mediates replicative crisis. Nature 614, 767–773. 10.1038/s41586-023-05710-8.

22. Hayashi, M.T., Cesare, A.J., Rivera, T., and Karlseder, J. (2015). Cell death during crisis is mediated by mitotic telomere deprotection. Nature 522, 492–496. 10.1038/nature14513.

23. Galluzzi, L., Vitale, I., Aaronson, S.A., Abrams, J.M., Adam, D., Agostinis, P., Alnemri, E.S., Altucci, L., Amelio, I., Andrews, D.W., et al. (2018). Molecular mechanisms of cell death: recommendations of the Nomenclature Committee on Cell Death 2018. Cell Death & Differentiation 25, 486–541. 10.1038/s41418-017-0012-4.

24. Masamsetti, V.P., Low, R.R.J., Mak, K.S., O’Connor, A., Riffkin, C.D., Lamm, N., Crabbe, L., Karlseder, J., Huang, D.C.S., Hayashi, M.T., and Cesare, A.J. (2019). Replication stress induces mitotic death through parallel pathways regulated by WAPL and telomere deprotection. Nature communications 10, 4224. 10.1038/s41467-019-12255-w.

25. Hayashi, M.T., Cesare, A.J., Fitzpatrick, J.A., Lazzerini-Denchi, E., and Karlseder, J. (2012). A telomere-dependent DNA damage checkpoint induced by prolonged mitotic arrest. Nature structural & molecular biology 19, 387–394. 10.1038/nsmb.2245.

26. Hain, K.O., Colin, D.J., Rastogi, S., Allan, L.A., and Clarke, P.R. (2016). Prolonged mitotic arrest induces a caspase-dependent DNA damage response at telomeres that determines cell survival. Scientific reports 6, 26766. 10.1038/srep26766.

27. Orth, J.D., Loewer, A., Lahav, G., and Mitchison, T.J. (2012). Prolonged mitotic arrest triggers partial activation of apoptosis, resulting in DNA damage and p53 induction. Molecular biology of the cell 23, 567–576. 10.1091/mbc.E11-09-0781.

28. Takai, H., Smogorzewska, A., and de Lange, T. (2003). DNA damage foci at dysfunctional telomeres. Curr Biol 13, 1549–1556.

29. Cesare, A.J., Hayashi, M.T., Crabbe, L., and Karlseder, J. (2013). The telomere deprotection response is functionally distinct from the genomic DNA damage response. Molecular cell 51, 141–155. 10.1016/j.molcel.2013.06.006.

30. Denchi, E.L., and de Lange, T. (2007). Protection of telomeres through independent control of ATM and ATR by TRF2 and POT1. Nature 448, 1068–1071.

31. Fouche, N., Cesare, A.J., Willcox, S., Ozgur, S., Compton, S.A., and Griffith, J.D. (2006). The basic domain of TRF2 directs binding to DNA junctions irrespective of the presence of TTAGGG repeats. The Journal of biological chemistry 281, 37486–37495. 10.1074/jbc.M608778200.

32. Schmutz, I., Timashev, L., Xie, W., Patel, D.J., and de Lange, T. (2017). TRF2 binds branched DNA to safeguard telomere integrity. Nature structural & molecular biology 24, 734–742. 10.1038/nsmb.3451.

33. Wang, R.C., Smogorzewska, A., and de Lange, T. (2004). Homologous recombination generates T-loop-sized deletions at human telomeres. Cell 119, 355–368.

34. Poulet, A., Buisson, R., Faivre-Moskalenko, C., Koelblen, M., Amiard, S., Montel, F., Cuesta-Lopez, S., Bornet, O., Guerlesquin, F., Godet, T., et al. (2009). TRF2 promotes, remodels and protects telomeric Holliday junctions. The EMBO journal 28, 641–651.

35. Carmena, M., Wheelock, M., Funabiki, H., and Earnshaw, W.C. (2012). The chromosomal passenger complex (CPC): from easy rider to the godfather of mitosis. Nature reviews 13, 789–803. 10.1038/nrm3474.

36. Bizard, A.H., and Hickson, I.D. (2014). The dissolution of double Holliday junctions. Cold Spring Harb Perspect Biol 6, a016477. 10.1101/cshperspect.a016477.

37. Hung, V., Udeshi, N.D., Lam, S.S., Loh, K.H., Cox, K.J., Pedram, K., Carr, S.A., and Ting, A.Y. (2016). Spatially resolved proteomic mapping in living cells with the engineered peroxidase APEX2. Nature protocols 11, 456–475. 10.1038/nprot.2016.018.

38. Kops, G.J., and Shah, J.V. (2012). Connecting up and clearing out: how kinetochore attachment silences the spindle assembly checkpoint. Chromosoma 121, 509–525. 10.1007/s00412-012-0378-5.

39. Sarek, G., Kotsantis, P., Ruis, P., Van Ly, D., Margalef, P., Borel, V., Zheng, X.F., Flynn, H.R., Snijders, A.P., Chowdhury, D., et al. (2019). CDK phosphorylation of TRF2 controls t-loop dynamics during the cell cycle. Nature 575, 523–527. 10.1038/s41586-019-1744-8.

40. Zimmermann, M., Kibe, T., Kabir, S., and de Lange, T. (2014). TRF1 negotiates TTAGGG repeat-associated replication problems by recruiting the BLM helicase and the TPP1/POT1 repressor of ATR signaling. Genes & development 28, 2477–2491. 10.1101/gad.251611.114.

41. Cesare, A.J., Heaphy, C.M., and O’Sullivan, R.J. (2015). Visualization of Telomere Integrity and Function In Vitro and In Vivo Using Immunofluorescence Techniques. Current protocols in cytometry / editorial board, J. Paul Robinson, managing editor … [et al 73, 12 40 11–12 40 31. 10.1002/0471142956.cy1240s73.

42. Meraldi, P., Honda, R., and Nigg, E.A. (2004). Aurora kinases link chromosome segregation and cell division to cancer susceptibility. Current opinion in genetics & development 14, 29–36. 10.1016/j.gde.2003.11.006.

43. Siemeister, G., Mengel, A., Fernandez-Montalvan, A.E., Bone, W., Schroder, J., Zitzmann-Kolbe, S., Briem, H., Prechtl, S., Holton, S.J., Monning, U., et al. (2019). Inhibition of BUB1 Kinase by BAY 1816032 Sensitizes Tumor Cells toward Taxanes, ATR, and PARP Inhibitors In Vitro and In Vivo. Clin Cancer Res 25, 1404–1414. 10.1158/1078-0432.CCR-18-0628.

44. Koltun, E.S., Tsuhako, A.L., Brown, D.S., Aay, N., Arcalas, A., Chan, V., Du, H., Engst, S., Ferguson, K., Franzini, M., et al. (2012). Discovery of XL413, a potent and selective CDC7 inhibitor. Bioorganic & medicinal chemistry letters 22, 3727–3731. 10.1016/j.bmcl.2012.04.024.

45. McKerlie, M., Lin, S., and Zhu, X.D. (2012). ATM regulates proteasome-dependent subnuclear localization of TRF1, which is important for telomere maintenance. Nucleic acids research 40, 3975–3989. 10.1093/nar/gks035.

46. Yaffe, M.B., Rittinger, K., Volinia, S., Caron, P.R., Aitken, A., Leffers, H., Gamblin, S.J., Smerdon, S.J., and Cantley, L.C. (1997). The structural basis for 14-3-3:phosphopeptide binding specificity. Cell 91, 961–971. 10.1016/s0092-8674(00)80487-0.

47. Neff, N.F., Ellis, N.A., Ye, T.Z., Noonan, J., Huang, K., Sanz, M., and Proytcheva, M. (1999). The DNA helicase activity of BLM is necessary for the correction of the genomic instability of bloom syndrome cells. Molecular biology of the cell 10, 665–676. 10.1091/mbc.10.3.665.

48. Machwe, A., Karale, R., Xu, X., Liu, Y., and Orren, D.K. (2011). The Werner and Bloom syndrome proteins help resolve replication blockage by converting (regressed) holliday junctions to functional replication forks. Biochemistry 50, 6774–6788. 10.1021/bi2001054.

49. Wu, L., Chan, K.L., Ralf, C., Bernstein, D.A., Garcia, P.L., Bohr, V.A., Vindigni, A., Janscak, P., Keck, J.L., and Hickson, I.D. (2005). The HRDC domain of BLM is required for the dissolution of double Holliday junctions. The EMBO journal 24, 2679–2687. 10.1038/sj.emboj.7600740.

50. Blackford, A.N., Nieminuszczy, J., Schwab, R.A., Galanty, Y., Jackson, S.P., and Niedzwiedz, W. (2015). TopBP1 interacts with BLM to maintain genome stability but is dispensable for preventing BLM degradation. Molecular cell 57, 1133–1141. 10.1016/j.molcel.2015.02.012.

51. Goulaouic, H., Roulon, T., Flamand, O., Grondard, L., Lavelle, F., and Riou, J.F. (1999). Purification and characterization of human DNA topoisomerase IIIalpha. Nucleic acids research 27, 2443–2450. 10.1093/nar/27.12.2443.

52. Suhasini, A.N., Rawtani, N.A., Wu, Y., Sommers, J.A., Sharma, S., Mosedale, G., North, P.S., Cantor, S.B., Hickson, I.D., and Brosh, R.M., Jr. (2011). Interaction between the helicases genetically linked to Fanconi anemia group J and Bloom’s syndrome. The EMBO journal 30, 692–705. 10.1038/emboj.2010.362.

53. Ohishi, T., Muramatsu, Y., Yoshida, H., and Seimiya, H. (2014). TRF1 ensures the centromeric function of Aurora-B and proper chromosome segregation. Molecular and cellular biology 34, 2464–2478. 10.1128/MCB.00161-14.

54. McKerlie, M., and Zhu, X.D. (2011). Cyclin B-dependent kinase 1 regulates human TRF1 to modulate the resolution of sister telomeres. Nature communications 2, 371. 10.1038/ncomms1372.

55. Chan, F.L., Vinod, B., Novy, K., Schittenhelm, R.B., Huang, C., Udugama, M., Nunez-Iglesias, J., Lin, J.I., Hii, L., Chan, J., et al. (2017). Aurora Kinase B, a novel regulator of TERF1 binding and telomeric integrity. Nucleic acids research 45, 12340–12353. 10.1093/nar/gkx904.

56. Scully, R., Panday, A., Elango, R., and Willis, N.A. (2019). DNA double-strand break repair-pathway choice in somatic mammalian cells. Nat. Rev. Mol. Cell Biol. 20, 698–714. 10.1038/s41580-019-0152-0.

57. Wyatt, H.D., Sarbajna, S., Matos, J., and West, S.C. (2013). Coordinated actions of SLX1-SLX4 and MUS81-EME1 for Holliday junction resolution in human cells. Molecular cell 52, 234–247. 10.1016/j.molcel.2013.08.035.

58. Ip, S.C., Rass, U., Blanco, M.G., Flynn, H.R., Skehel, J.M., and West, S.C. (2008). Identification of Holliday junction resolvases from humans and yeast. Nature 456, 357–361. 10.1038/nature07470.

59. West, S.C., Blanco, M.G., Chan, Y.W., Matos, J., Sarbajna, S., and Wyatt, H.D. (2015). Resolution of Recombination Intermediates: Mechanisms and Regulation. Cold Spring Harbor symposia on quantitative biology 80, 103–109. 10.1101/sqb.2015.80.027649.

60. Balbo Pogliano, C., Ceppi, I., Giovannini, S., Petroulaki, V., Palmer, N., Uliana, F., Gatti, M., Kasaciunaite, K., Freire, R., Seidel, R., et al. (2022). The CDK1-TOPBP1-PLK1 axis regulates the Bloom’s syndrome helicase BLM to suppress crossover recombination in somatic cells. Sci Adv 8, eabk0221. 10.1126/sciadv.abk0221.

61. Pickett, H.A., Cesare, A.J., Johnston, R.L., Neumann, A.A., and Reddel, R.R. (2009). Control of telomere length by a trimming mechanism that involves generation of t-circles. The EMBO journal 28, 799–809. 10.1038/emboj.2009.42.

62. Barber, L.J., Youds, J.L., Ward, J.D., McIlwraith, M.J., O’Neil, N.J., Petalcorin, M.I., Martin, J.S., Collis, S.J., Cantor, S.B., Auclair, M., et al. (2008). RTEL1 maintains genomic stability by suppressing homologous recombination. Cell 135, 261–271. 10.1016/j.cell.2008.08.016.

63. Vannier, J.B., Pavicic-Kaltenbrunner, V., Petalcorin, M.I., Ding, H., and Boulton, S.J. (2012). RTEL1 dismantles T loops and counteracts telomeric G4-DNA to maintain telomere integrity. Cell 149, 795–806. 10.1016/j.cell.2012.03.030.

64. Orthwein, A., Fradet-Turcotte, A., Noordermeer, S.M., Canny, M.D., Brun, C.M., Strecker, J., Escribano-Diaz, C., and Durocher, D. (2014). Mitosis inhibits DNA double-strand break repair to guard against telomere fusions. Science (New York, N.Y 344, 189–193. 10.1126/science.1248024.

65. Romero-Zamora, D., and Hayashi, M.T. (2023). A non-catalytic N-terminus domain of WRN prevents mitotic telomere deprotection. Scientific reports 13, 645. 10.1038/s41598-023-27598-0.

66. Timashev, L.A., and De Lange, T. (2020). Characterization of t-loop formation by TRF2. Nucleus-Phila 11, 164–177. 10.1080/19491034.2020.1783782.

67. Lam, S.S., Martell, J.D., Kamer, K.J., Deerinck, T.J., Ellisman, M.H., Mootha, V.K., and Ting, A.Y. (2015). Directed evolution of APEX2 for electron microscopy and proximity labeling. Nature methods 12, 51–54. 10.1038/nmeth.3179.

68. Garcia-Exposito, L., Bournique, E., Bergoglio, V., Bose, A., Barroso-Gonzalez, J., Zhang, S., Roncaioli, J.L., Lee, M., Wallace, C.T., Watkins, S.C., et al. (2016). Proteomic Profiling Reveals a Specific Role for Translesion DNA Polymerase eta in the Alternative Lengthening of Telomeres. Cell reports 17, 1858–1871. 10.1016/j.celrep.2016.10.048.

69. Barefield, C., and Karlseder, J. (2012). The BLM helicase contributes to telomere maintenance through processing of late-replicating intermediate structures. Nucleic acids research 40, 7358–7367. 10.1093/nar/gks407.

70. Bosco, N., and de Lange, T. (2012). A TRF1-controlled common fragile site containing interstitial telomeric sequences. Chromosoma 121, 465–474. 10.1007/s00412-012-0377-6.

71. Hsieh, M.Y., Fan, J.R., Chang, H.W., Chen, H.C., Shen, T.L., Teng, S.C., Yeh, Y.H., and Li, T.K. (2014). DNA topoisomerase III alpha regulates p53-mediated tumor suppression. Clin Cancer Res 20, 1489–1501. 10.1158/1078-0432.CCR-13-1997.

72. Chen, E.Y., Tan, C.M., Kou, Y., Duan, Q., Wang, Z., Meirelles, G.V., Clark, N.R., and Ma’ayan, A. (2013). Enrichr: interactive and collaborative HTML5 gene list enrichment analysis tool. BMC bioinformatics 14, 128. 10.1186/1471-2105-14-128.

73. Cesare, A.J., Kaul, Z., Cohen, S.B., Napier, C.E., Pickett, H.A., Neumann, A.A., and Reddel, R.R. (2009). Spontaneous occurrence of telomeric DNA damage response in the absence of chromosome fusions. Nature structural & molecular biology 16, 1244–1251. 10.1038/nsmb.1725.

